# Amendable decisions in living systems

**DOI:** 10.1101/2025.06.03.657747

**Authors:** Izaak Neri, Simone Pigolotti

## Abstract

A distinct feature of living systems is their capacity to take decisions based on uncertain environmental signals. Examples span from the microscopic scale of cell differentiation guided by the concentration of a morphogen, to complex choices made by animals and humans. The current paradigm in decision theory assumes that decisions, once taken, cannot be revoked. However, living systems often amend their decisions if new evidence favors an alternative hypothesis. In this paper, we characterize the optimal strategy for such amendable decisions. We find that, unlike irrevocable decisions, optimal amendable decisions can be made in a finite average time with zero error probability. Our theory successfully predicts the outcome of a visual experiment involving human participants and the accuracy of cell-fate decisions in early development. We also present possible implementations of amendable decision-making by means of either biochemical reactions or transcription regulation. Our study reveals that amendments lead to a substantial advantage in decision making, that is likely to be widely exploited by living beings.

---

Living systems are able to make appropriate decisions based on signals received from their environment. Examples of evidence-based decision-making can be found in animal behavior [1], such as in mate choice [2], foraging [3, 4], and prey and predator detection [5]. Decision tasks have been studied in laboratory experiments, where performance can be tested under controlled conditions [1, 4, 6, 7]. In the wild, by contrast, deciding rapidly and appropriately is often a matter of life and death. For this reason, living systems are often argued to employ optimal decision strategies [8–10].

We focus on decision processes in which a living observer has in mind different, mutually exclusive hypotheses about their environment. The observer accumulates information about the environment, and stores it in an internal representation called the evidence. At some point in time, the observer decides, based on the evidence, which of the hypotheses they believe to be correct. Literature on biological decision making has predominantly focused on decisions that cannot be revoked. In this setup, the optimal strategy has been identified by Wald as the sequential probability ratio test [11–13]. Here, the “optimal strategy” is defined as the fastest on average in reaching a decision for a prescribed error probability (the probability of committing to the wrong hypothesis). In the sequential probability ratio test, the evidence is constructed as the log likelihood ratio of the two hypotheses, and a decision is made as soon as the evidence reaches either a positive or a negative threshold. Neural network models support that the cortico-basal ganglia circuit in mammalian brains indeed implements a sequential probability ratio test [14, 15].

The sequential probability ratio test is characterized by speed-accuracy tradeoffs [1, 16, 17]. For example, the average decision time increases as the similarity between the hypotheses is increased. Additionally, the average decision time increases as the error probability is decreased. Speed-accuracy tradeoffs have been confirmed in behavioral experiments on olfactory [7], visual [3, 6], and tactile [18] tasks performed by animals [3, 7, 18, 19] and humans [6]; see [1, 17] for reviews.

Certain fast and accurate cellular processes, such as the expression of genes in response to morphogen signals [8, 20–2] T-cell activation in response to foreign ligands [24], and stem cell differentiation [25] can be conceptualized as decisions as well [26–28] It has been proposed that cells, like brain networks, make decisions based on the sequential probability ratio test, or at least on a biochemical approximation of this algorithm [23, 28, 29], although stringent experimental tests of this idea are still lacking.

Irrevocable decisions are often unnatural for living beings. Perceptual studies have shown that humans often change their mind after having perceived a stimulus and taken a decision, thereby reducing their error probability [30–32] It is also debatable whether cellular decisions are strictly irrevocable. Subcellular regulatory systems responsible for decisions can be conceptualized as noisy high-dimensional dynamical systems settling into an attractor [33–35] Because of noise, these systems gradually commit to a particular outcome, rather than undergo a sudden irreversible transition. In addition, many cell-fate decisions can be amended through de-differentiation or trans-differentiation [36]. For example, hepatocytes convert to biliary cells after liver injury over a period of weeks through stepwise changes in gene expression, and cells that have not completed the transition revert to their original identity once the stimulus is removed [37, 38].

In this paper, we present a theory for decision processes in which, after decisions, observers maintain their right to change their mind, should additional evidence overwhelmingly favor the alternative hypothesis. We identify the optimal strategy in this scenario. In stark contrast with irrevocable decisions, this optimal amendable strategy is characterized by a vanishing error probability in a finite average decision time. We apply our theory to a human perception study, finding support for our theoretical predictions. We then apply our theory to the microscopic world of cell-fate decisions. We show that amendable decisions can be used by cell receptors to implement a concentration test, leading to a speed advantage over irrevocable decisions. We also apply this idea to the Bicoid-Hunchback system, a paradigmatic system for cell-fate decisions in development. We show that the amendable model successfully predicts the experimental decision statistics with one fitting parameter, the Hunch-back boundary position, where the expression probability crosses 1/2. We conclude by presenting possible implementations of amendable decisions in transcription regulation.

The paper is organized as follows. In Section I, we formally define irrevocable and amendable decision processes. In Section II, we apply the theory to an observation signal described by a drift-diffusion process. In Section III, we study the optimal evidence process for amendable decisions. In Section IV, we compare irrevocable and amendable decisions in a visual perception experiment. In Section V, we study optimal amendable decisions by a molecular receptor sensing concentrations of incoming molecules. In Section VI, we propose a biochemical circuit implementing the amendable concentration test. In Section VII, we compare optimal irrevocable and amendable decision models for the Bicoid-Hunchback system. In Section VIII, we present a model for amendable decisions in transcription regulation which is applicable to the Bicoid-Hunchback system, as well as to other genes regulated by a transcription factor. Section IX is devoted to conclusions and perspectives.

## I. IRREVOCABLE VERSUS AMENDABLE DECISIONS

We focus on decisions involving two hypotheses, + and −, on a certain phenomenon. An observer acquires information about this phenomenon by perceiving a time-dependent signal *X* (*t*), see Fig. 1a. This signal is noisy, and its statistics depend on which of the two hypotheses holds. The observer monitors the signal and stores the information contained in it into a time-dependent variable *E* (*t*), called the evidence. This variable is constructed so that positive and negative values of *E* represent accumulated evidence supporting hypotheses + and −, respectively. The decision is a binary variable *D* ∈ {+ −}, that depends on the evidence and is considered correct if it matches the hypothesis.

**FIG. 1.**
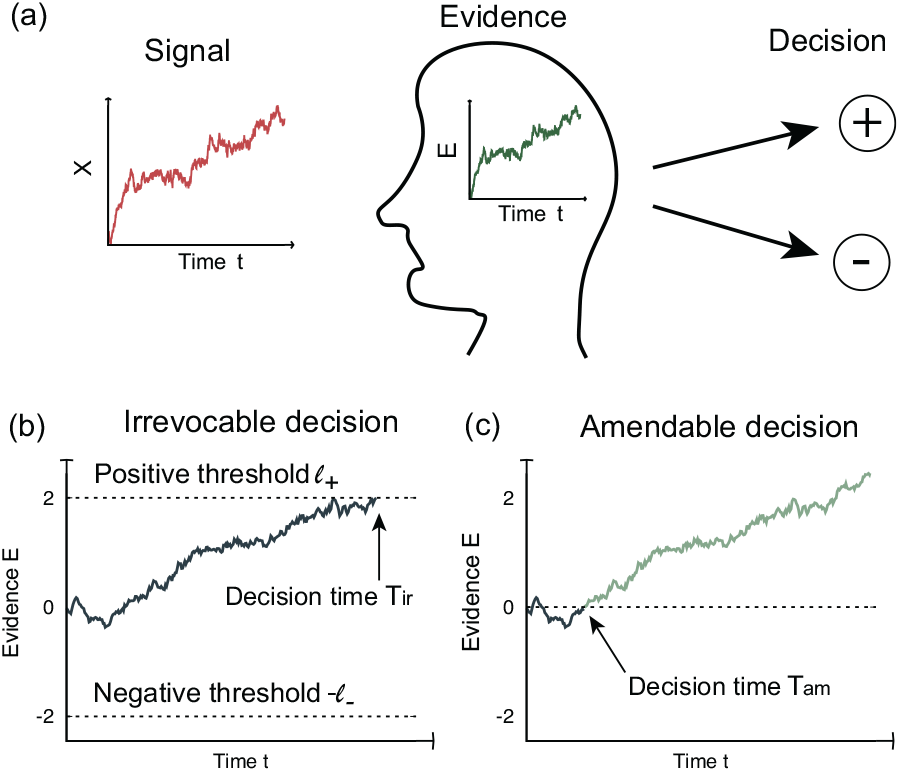
Illustration of the difference between irrevocable and amendable decisions. (a) Evidence-based decision making. An environmental signal is observed by a living system that stores the information contained in it in a process called the evidence. After having accumulated enough information, the observer makes a binary decision. (b) Irrevocable decision making. The observer commits to the + or - hypothesis as soon as the evidence crosses a positive or negative threshold, respectively. This commitment is irrevocable. (c) Amendable decision making. In this case, the observer is in favor of the + or − hypothesis whenever *E* (*t*) is greater or smaller than zero, respectively. A final decision time can be identified a posteriori as the moment when the evidence process crosses zero for the last time.

We define decisions in a different way for irrevocable and amendable processes. In an irrevocable decision, the observer commits to the hypothesis + (or−) as soon as the collected evidence crosses either a positive threshold −*l*_+_ or a negative threshold −*l*_−_, see Fig. 1b. This commitment is final. This idea is at the basis of Wald’s sequential test [11, 12], in which the evidence is constructed as the logarithm of the likelihood ratio of the two hypotheses. The decision time *T*_ir_ is the time when either of the two thresholds is first crossed. Properties of irrevocable decision processes can be calculated using the theory of first-passage times [39]. Choosing larger thresholds amounts to safer decisions, i.e., a lower probability of committing to the wrong hypothesis. However, this comes at a cost of a longer decision time.

In an amendable decision process, observers can alter their choice, so that their decision is only tentative at any time during the observation. We call *d* (*t*) ∈ {+, −}, the tentative decision at time *t*. The natural strategy is to set *d*(*t*) equal to the sign of *E*(*t*). In practice, the evidence *E* (*t*) can change sign several times, leading to multiple amendments to the tentative decision. However, if the observer keeps gathering evidence and time is unlimited, the evidence *E* (*t*) will eventually drift so far from the origin that a reversal becomes vanishingly unlikely (see Fig. 1). We then define the final decision *D* as the large-time limit of *d t* . Alternatively, one could consider scenarios in which observers have a finite available time (*t*_f_) to reach a decision, in which case we define the final decision *D* as *d* (*t*_f_). In both cases, we identify the decision time *T*_am_ a posteriori as the time when the evidence process crosses zero for the last time, see Fig. 1c.

## II. ERROR-FREE AMENDABLE DECISIONS IN THE DRIFT-DIFFUSION MODEL

We consider an observation signal *X* (*t*) whose statistics are described by a drift-diffusion model

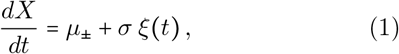

where *µ*_+_ and *µ*_−_ are the drift coefficients corresponding to the two hypotheses and *ξ*(*t*) is white noise. Without loss of generality, we assume *µ*_+_ > *µ*_−_. We express the dynamics of the evidence process by

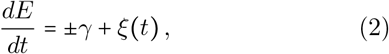

where the plus/minus sign in the right hand side depends on which of the two hypotheses hold, and *γ* is the average rate of evidence acquisition. In Eq. (2), we have set the amplitude of the noise equal to one by an appropriate choice of the units of *E*. We have also assumed that changing hypothesis simply changes the sign of the drift term. This assumption is justified in two cases: when the observation process is itself symmetric, i.e., when *µ*_−_ = −*µ*_+_; or if the evidence process is constructed as the log-likelihood ratio process *E* (*t*) ∝ ln *P* (*X* (*t*) ∣+)/*P* (*X* (*t*)∣−) (see Appendix C). Here, *P* (*X*(*t*)∣+) and *P* (*X*(*t*)∣−) are the probability distributions of *X* (*t*) given the positive and negative hypothesis, respectively. More general evidence processes, for which the magnitude of the rate of evidence acquisition depends on the hypothesis, are discussed in Appendices A and B.

For an irrevocable decision, the optimal strategy is given by the Wald test, for which the evidence process equals the logarithm of the likelihood ratio. This choice is optimal in the sense of minimizing the average irrevocable decision time at a given error probability (see Ref. [40] or Appendix D). The optimal evidence obeys Eq. (2) with 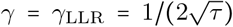, where we have introduced the characteristic time *τ* = *σ*^2^ /(*µ*_+_ − *µ*_−_)^2^ (see Appendix C). We focus on the case of symmetric barriers *l*_+_ *= l*_− =_ *l*, where the probabilities of false positives and false negatives are equal.

We compare statistics of the decision time and error probability for irrevocable and amendable decisions based on the log-likelihood evidence process. In the irrevocable case, the average decision time depends on the choice of the thresholds, and thus on how the observer trades off speed and error probability, see Fig. 2a. In particular, for small error probability *η*, the average decision time scales as ⟨*T*_ir_ ⟩ ∼ − log(*η*). Here and in the following, we denote by ⟨… ⟩ an average over the observation process and over the two hypotheses. In contrast, amendable decisions are free of errors (*η* = 0) and are reached in a finite average decision time ⟨*T*_am_⟩ = 4*τ* . Amendable decisions are faster than irrevocable decisions on average, unless the observer is willing to accept an error probability larger than a critical value *η*_*c*_ ≈ 0.08), see Fig. 2a. While the distribution *p*_ir_ *t* of irrevocable decision times presents a peak at an intermediate time, the distribution of amendable decision times

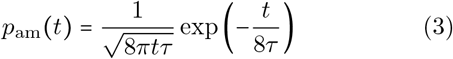

decreases monotonically, see Fig. 2b.

**FIG. 2.**
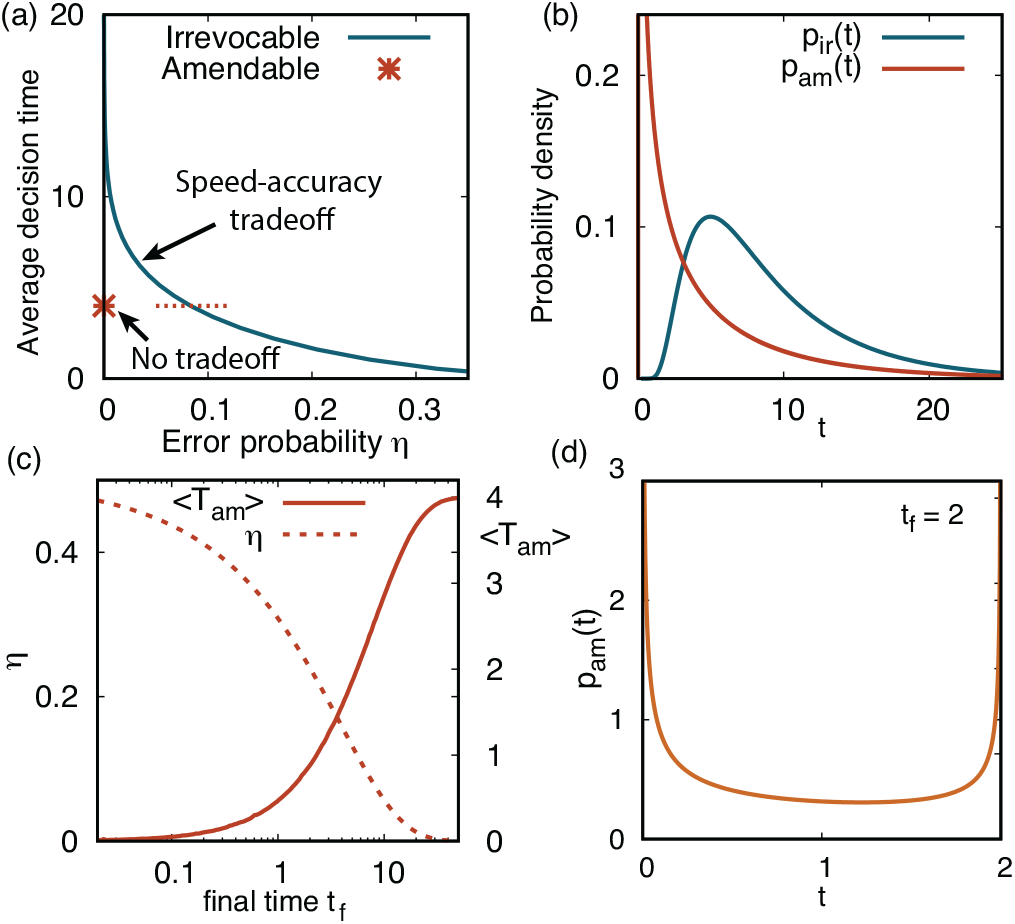
Comparison between irrevocable and amendable decisions in the drift diffusion process. Both decisions use the same evidence process *E* (*t*)governed by Eq. (2), with 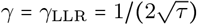 and units of time *τ* = 1. (a) Comparison faster for *η* > *η*_*c*_ ≈ 0.083. (b) Decision time distributions for of average decision times. In the irrevocable case, the decision time depends on the error probability *η*, while amendable decisions are always error-free (*η* = 0). Irrevocable decisions are amendable and irrevocable decisions for infinitely large observation time *t*_f_ . In the irrevocable case, we have set *η* = 0.01. (c) Average decision time and error probability for amendable decisions within a finite observation time, *t* ∈ [0], *t*_f_ . (d) Distribution of amendable decision times for a finite observation time. The results shown in this figure are derived in Appendices A and B.

So far, we have considered cases in which the time available to the observer is unlimited. However, in practical situations, the observer might be forced to make a call within a certain maximum time *t*_f_ . In such settings, amendable decisions are characterized by a finite error probability, see Fig. 2c. For small *t*_f_, the distribution of decision times presents a bimodal shape, see Fig. 2d. The second peak close to *t*_f_ is due to the conditioning of a decision before the maximum time: a transition via the state *E =* 0 has an increased chance of being the last one if it occurs just before *t*_*f*_ .

## III. OPTIMAL EVIDENCE PROCESS FOR AMENDABLE DECISIONS

We have compared amendable and irrevocable decisions in which the evidence is chosen as the log-likelihood ratio, a strategy that is optimal for irrevocable decisions [40]. Here, we scrutinize whether the log likelihood ratio is also optimal for amendable decisions. As in Section II, we consider observations described by the drift-diffusion process, Eq. (1). Using that *X* (*t*) is a time-homogeneous Markov process and assuming that the evidence is an additive functional of *X* (*t*), the general evidence process takes the form

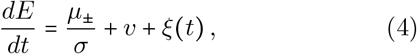

see Appendix C. We have set the noise amplitude in Eq. (4) to one using that the absolute scale of the evidence process is irrelevant for amendable decisions. Hence, the two parameters that specify the evidence process are the constant *v* appearing in Eq. (4) and the initial condition *E*_0_. The log-likelihood process corresponds to choosing *v =* 0 and *E*_0_ = 0.

We consider the class of evidence processes defined in Eq. (4) and minimize the mean amendable decision time ⟨*T*_am_⟩ with respect to *v* and *E*_0_, subject to the constraint that the error probability vanishes (see Appendix E). We find that the optimal evidence process is characterized by non-trivial values of *v* and *E*_0_. The log-likelihood ratio only approximates the optimal process, see Fig. 3 and Appendix E. Nevertheless, the difference of performance between the log-likelihood process and the optimal process is quite small: for example, in the case of uniform prior, the average decision time for the optimal process is ⟨*T*_am_⟩ ≈ 3.9949 *τ* compared with ⟨*T*_am_⟩ ≈ 4*τ* for the log-likelihood process, with *τ = σ*^2^ /(*µ*_+_ − *µ*_−_)^2^ the characteristic time scale. We have additionally found that the log-likelihood process is optimal for amendable decisions in terms of minimizing the maximum average decision time between the two hypotheses, see Appendix F.

**FIG. 3.**
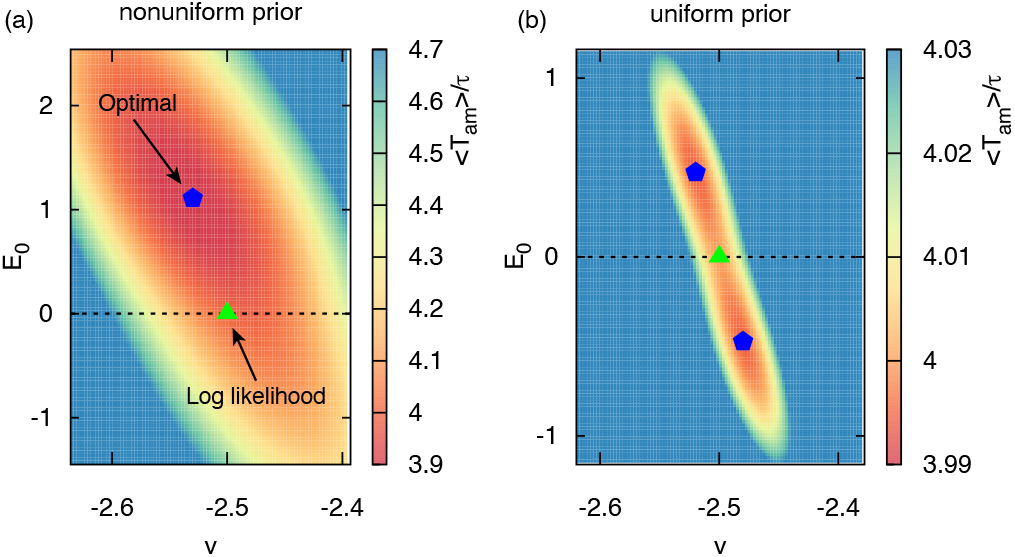
Average amendable decision time ⟨*T*_am_⟩/*τ* as a function of the parameters *E*_0_ and *v* of the evidence process, see Eq. (4). Panel (a): nonuniform prior, *π*_+_ = 1_−_ − *π* = 0.55. the probabilities that *H* = + and *H* = − are true, respectively. Panel(b): uniform prior, *π*_+_ = *π*_−_ = 0.5, where *π*_+_ and *π*_−_ are Other model parameters are *µ*_+_ = 3, *µ*_−_ = 2, and *σ* = 1. See Appendix E for details.

In summary, the log-likelihood evidence process performs closely to the optimal one, and additionally it minimizes the maximum of the two average decision times conditioned on the hypotheses. For these reasons, evidence-based decision making based on the log-likelihood ratio will be the object of our study for the rest of the paper. This choice has also the advantage of allowing for a more direct comparison between irrevocable and amendable decisions.

## IV. AMENDABLE VS. IRREVOCABLE DECISIONS IN A VISUAL PERCEPTION EXPERIMENT

To examine how amendable decision theory applies in the real world, we performed a human visual perception experiment. Participants were asked to observe the motion of a randomly moving dot on a screen, and assess whether the motion was biased towards the left or right direction by pressing a key, see Fig. 4a. In the irrevocable version of the task, pressing a key terminated the game. In the amendable version, participants could alter their decision an arbitrary number of times, until a time limit *t*_f_ = 10 s. At the initial time, the tentative decision *d*(*t*) was randomly initialized to + or − with equal probabilities. Both in the irrevocable and amendable version, participants played four different versions of the games of increasing difficulty, i.e., reduced bias. Participants were instructed to decide as fast as possible, but without making too many mistakes. Additional details on the experimental protocols are presented in Appendix G.

**FIG. 4.**
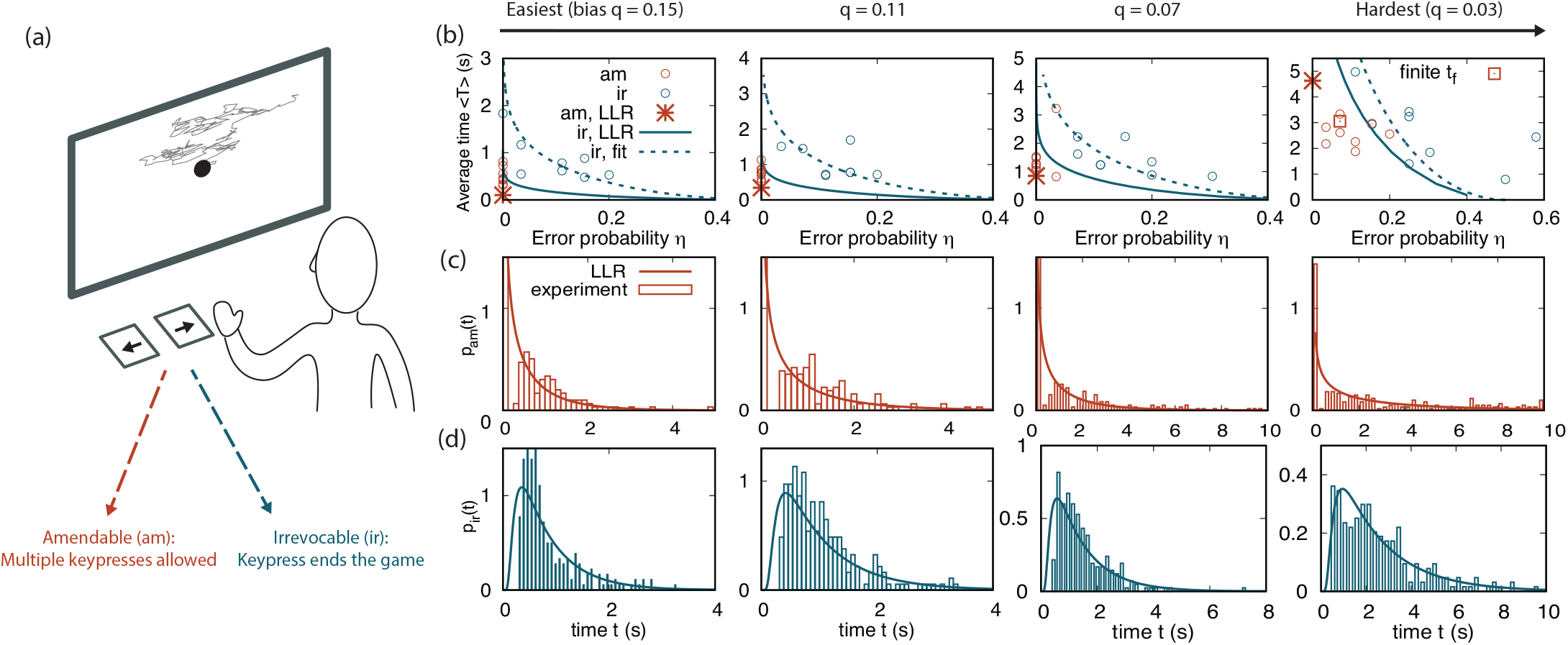
Amendable versus irrevocable decisions in a visual perception experiment. (a) Sketch of the experiment. (b) Speed-error tradeoffs for the four game difficulties. The game difficulty is determined by the bias parameter *q* (see Appendix G). Circles represent the error probabilities and average decision times for each individual player. In the irrevocable cases, average decision times and error probabilities are characterized by a tradeoff (Pearson correlation coefficients among participants *r* = {−0.68, −0.48, −0.61, −0.61} from easiest to hardest). The star and the continuous line mark the theoretical results for the log-likelihood ratio (LLR) evidence process in the amendable and irrevocable cases, respectively (see Fig. 2a). In the LLR case, the parameter *γ*_LLR_ is computed from the drift and diffusion of the moving dot (see Appendix H). The dashed lines represent the irrevocable strategy associated with a fitted evidence rate *γ* (see Appendix H). For *q* = 0.03, the square marks the finite-time error and average decision time for the log-likelihood case (see Appendix B and H). For all the other game difficulties, the finite-time amendable strategy is practically indistinguishable from the infinite-time one, and thus decisions are made with negligible error probability. (c) Empirical distributions of the amendable decision time, *p*_am_(*t*), compared with the theoretical distribution given by Eq. (3). (d) Empirical distributions of the irrevocable decision time, *p*_ir_(*t*), compared with the optimal distribution estimated from the empirical error probability (see Appendix H).

In keeping with the literature [6], we model evidence accumulation as a drift-diffusion process. We consider both a normative model [41] in which we set the evidence rate to the value corresponding to the log-likelihood ratio (*γ* = *γ*_LLR_) and a descriptive model in which we infer *γ* from the experimental data. The log-likelihood evidence rate is determined by the drift and diffusion coefficients of the moving dot. Therefore, the normative amendable decision model has no fitting parameters. In contrast, the normative irrevocable decision model includes the threshold *l* as a free parameter.

As predicted by the normative model, participants were able to make error-free amendable decisions, at least when the task was not too difficult (Fig. 4b). In the hardest version of the game, errors took place. This can be explained within the theory due to the finite duration of each trial (see Fig. 2c and Appendix B). In contrast, irrevocable decisions were error-prone for all game difficulties, and characterized by a speed-error tradeoff (Fig. 4b).

We now compare the empirical distributions of decision times with those of the normative drift-diffusion model. For easy games, the empirical distributions present a “gap” for short times (on the order of 0.1s) compared with the theoretical prediction, see Fig. 4c and Fig. 4d. This discrepancy is likely due to the finite response time of players. The tails of the distributions are instead in quantitative agreement with the theoretical ones. For harder games, we find an excellent agreement between the theory and the experiment for all times. This is consistent with the observation that the empirical drift is close to the normative one for these games, suggesting that players adopt a nearly optimal strategy. This result could be, again, due to the fact that harder games involve longer decision times, making individual reaction time less of a limiting factor. Empirical decision-time distributions for irrevocable games are qualitatively different than those for amendable ones, with a distinct peak at intermediate times for irrevocable decisions, as predicted by the theory (Fig. 4d).

For the descriptive model, we used the experimental data to estimate the evidence rate *γ* for each participant and each game, see Fig. 5 and Appendix H. In most cases, we found higher evidence rates in amendable than in irrevocable games (see Fig. 5), and therefore closer to the normative evidence rates. This suggests that amendable decision tasks capture the rate of acquired information with higher accuracy than irrevocable tasks. Furthermore, for each of the games, tradeoffs curves calculated for the evidence rate averaged over the participants are in good agreement with participants performance (dashed lines in Fig. 4b). An extended analysis of speed-error tradeoffs for individual participants is presented in Appendix I.

**FIG. 5.**
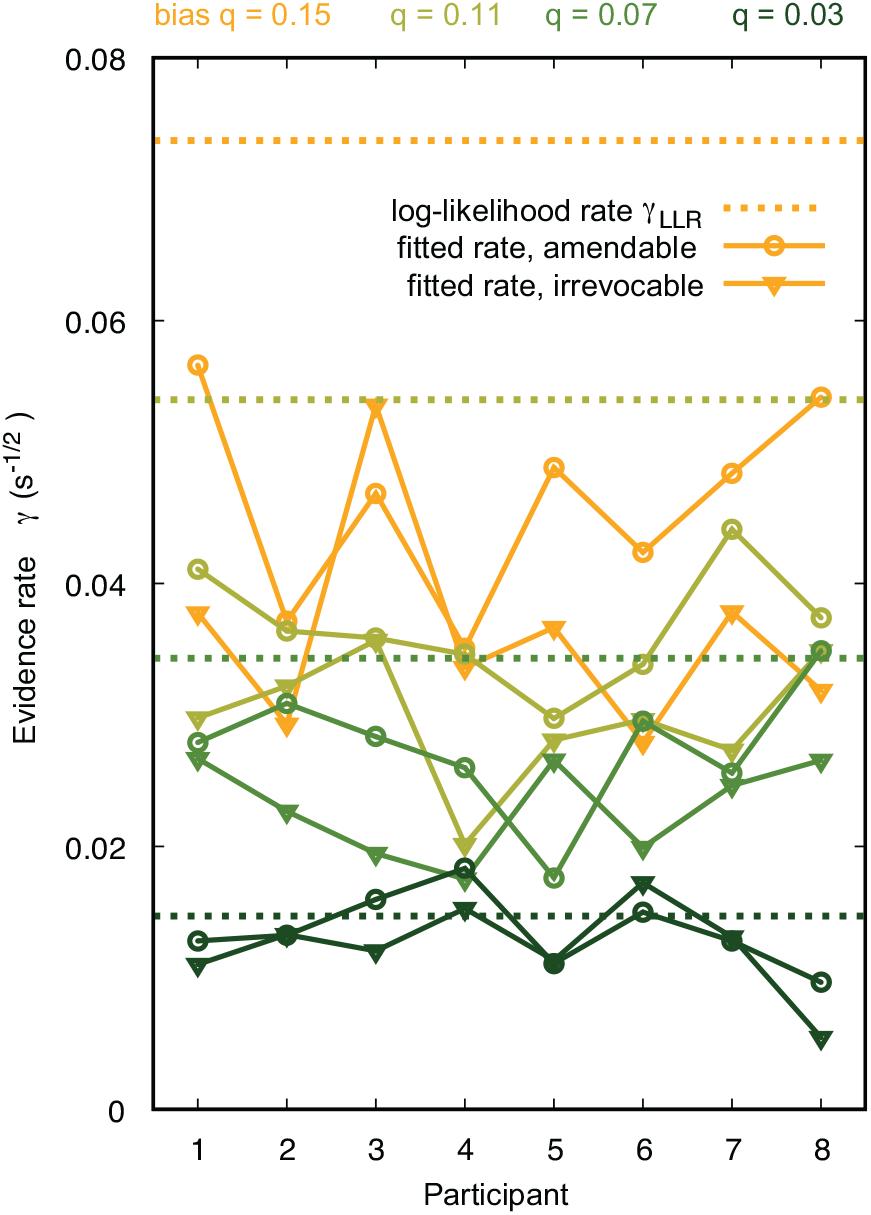
Estimated rates of evidence acquisition *γ* for the eight participants and the four game difficulties (markers) in the visual perception experiment of Fig. 4 are compared with the log-likelihood evidence rates 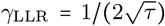 (dotted lines) with *τ* = Δ*t* /(16*q*^2^) (see Appendix H). As expected, most of the empirical data points lie below the corresponding LLR values (91% of data points, exceptions are participant 8 in the amendable game with *q* = 0.07, participants 4 and 6 for the irrevocable game with *q* = 0.03, and participants 3, 4 and 6 for the amendable game with *q* = 0.03). In most cases, participants displayed a higher evidence rate for amendable than irrevocable games (81% of data points, exceptions are participant 3 for *q* = 0.15, participant 5 for *q* = 0.07, and participants 2, 5, 6, and 7 for *q* = 0.03). For amendable games, the evidence rates are estimated from 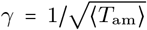, with ⟨*T*_am_⟩ the empirically measured value of the mean decision time for each game difficulty. For the hardest game *q* = 0.03, we used theoretical expressions for ⟨*T*_am_⟩ and *η* at finite time (see Appendix B). For irrevocable games, the evidence rates are estimated from the data as explained in Appendix H.

## V. AMENDABLE DECISIONS IN CELLULAR SENSING

Biological cells often need to accurately measure the concentration of certain molecules, for example, extracellular ligands or nuclear transcription factors, and make decisions depending on the outcome of the measurement. These decisions can be conceptualized within our framework, with the “observer” being the component of the cell responsible for detecting the molecules. In keeping with the literature [42, 43], we model the observer as a spherical body that absorbs molecules present in a surrounding fluid at a certain concentration *c*_∞_ far from itself, see Fig. 6a. We assume that the environment is at thermodynamic equilibrium, so that the stationary current of absorbed molecules is given by the Smoluchowski formula *j =* 4*πrDc*_∞_, where *r* is the radius of the observer and *D* the molecular diffusion coefficient [42]. The observer’s task is to distinguish between two values of the concentration, *c*_+_ and *c*_−_, with *c*_+_ > *c*_−_, corresponding to two different rates of incoming molecules, *j*_+_ and *j*_−_, respectively. The number of molecules *N* _±_ (*t*) absorbed up to time *t* represents our observation process. The precision parameter *ϵ* = (*c*_+_ − *c*_−_)/*c*_+_ quantifies the relative difference between the two concentrations, and therefore the difficulty of the discrimination task.

**FIG. 6.**
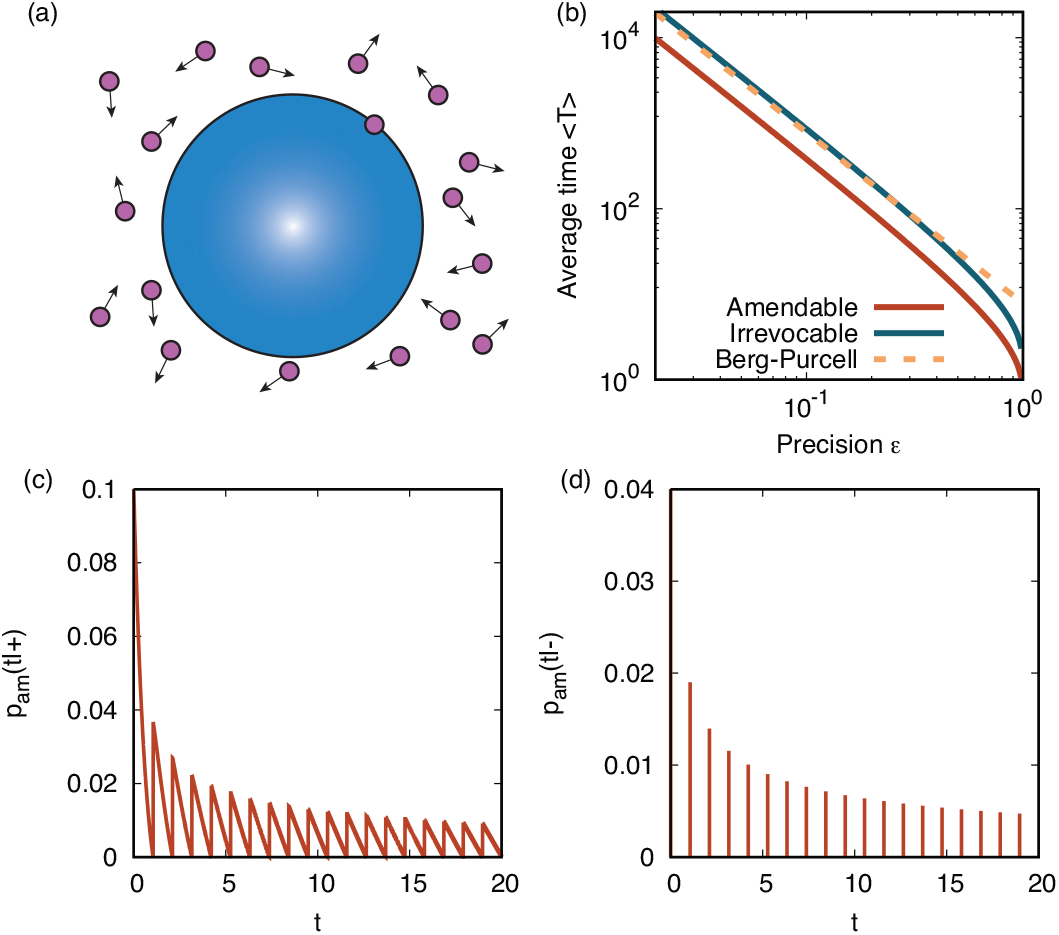
Decisions in molecular sensing. (a) A microscopic observer absorbs molecules from the environment attempting to detect their concentration. (b) Average decision times as a function of the precision *ϵ*. We have set τ = 1. The Berg-Purcell line is the estimate ⟨*T* ⟩ = (4*π*/(1.61))*ϵ*^−2^ for the mean decision time, that is based on the root mean square fractional error of the concentration, see Eq. (30) in [42]. For irrevocable decisions we have set *η* = 0.01. (c) Distribution of amendable decision times for the + hypothesis and *ϵ* = 0.1. (d) Same as (c), but for the − hypothesis. The distribution is discrete in this case (see Appendix K for analytical expressions).

In this case, the log likelihood ratio process is governed by the equation

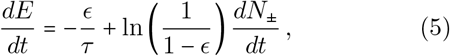

see Appendix K and [28]. Here, *τ* = 1 /*j*_+_ is the characteristic time scale of the dynamics.

We compare the average decision times for amendable decisions, irrevocable decisions, and the Berg-Purcell prediction [42] in Fig. 6b. In all three cases, the average decision time scales as *ϵ* ^−2^ (see Appendix K). Amendable decisions are error-free and about a factor two faster than the Berg-Purcell prediction. The average irrevocable decision time depends on the chosen error probability and it is larger than the amendable one for *η* < 0.083, approximately, as for the drift-diffusion model (see Appendix K). Thus, amendable decisions in molecular recognition based on the log-likelihood ratio process are error-free and faster than the optimal irrevocable decisions model, independently of the difficulty *ϵ* of the discrimination task.

Mathematical analyses of receptor models often approximate the input signal as a drift-diffusion process [28]. We derived the distribution of amendable decision times without resorting to such approximation, see Appendix K. The solution shows how the discrete nature of the information obtained from incoming molecules affects the statistics of the decision time. The distribution of decision times is discontinuous, and markedly different depending on whether the plus or minus hypothesis holds. In particular, for the positive hypothesis (higher concentration), the distribution presents a sawtooth shape (Fig. 6c). In contrast, for the negative hypothesis, the distribution is nonzero only over a discrete set of times (Fig. 6d). This difference is due to an asymmetry in the evidence dynamics embodied in Eq. (5): the evidence increases in discrete steps whenever the observer absorbs a molecule, while it declines continuously in time. In particular, in the negative hypothesis case, each of the discrete decision times corresponds to a given number of molecules absorbed before the decision (see Appendix K).

## VI. BIOCHEMICAL MODEL FOR AMENDABLE DECISIONS BY A RECEPTOR

We now study how an amendable concentration test can be implemented in a biochemical system. We develop a model in which both the evidence process *E*(*t*)and the decision process *d t* are expressed in terms of concentrations of molecules in a chemical reaction network. Our model is inspired by Ref. [28], that proposed an irrevocable decision model in which the evidence is encoded into two enzymes at concentrations *E*_1_ and *E*_2_. The first enzyme exponentially decays when the receptor is not bound to incoming molecules, while the second decays when molecules are bound. We follow the same idea, but in the limit of very large off-rates, in keeping with Section V. In this limit, *E*_1_ decays at a constant exponential rate and *E*_2_ decays in discrete steps at each binding event:

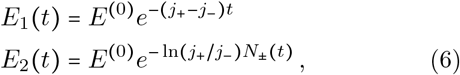

where *E*^(0)^ is the initial concentration, which we assumed identical for the two enzymes, and *N*_±_(*t*) is the cumulative number of received molecules, depending on the hypothesis. The two decay rates are such that

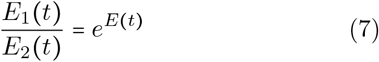

where *E* (*t*) is the log likelihood evidence process as given in Eq. (5). Equation (7) provides an invertible mapping between the evidence and the concentration ratio of the two enzymes.

We now move to the decision. A natural way to implement the decision is by means of a biochemical switch, i.e., a system capable of producing a sharp sigmoidal response as a function of an input. In particular, the model in Ref. [28] couples the two enzymes *E*_1_ and *E*_2_ with two downstream switches, specifically two Goldbeter-Koshland (GK) modules [44]. The kinetic rates of the two switches are such that their transitions occur when the evidence crosses the two absorbing thresholds *l*_+_ and −*l*_−_ in the Wald test. While in this approach the GK modules serve to represent irrevocable decisions, a GK module is actually a reversible switch. This property makes it particularly suitable for the amendable case.

We implement amendable decisions via a single GK module, see Fig. 7a. The GK module is described by the reactions

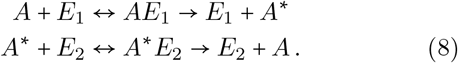

**FIG. 7.**
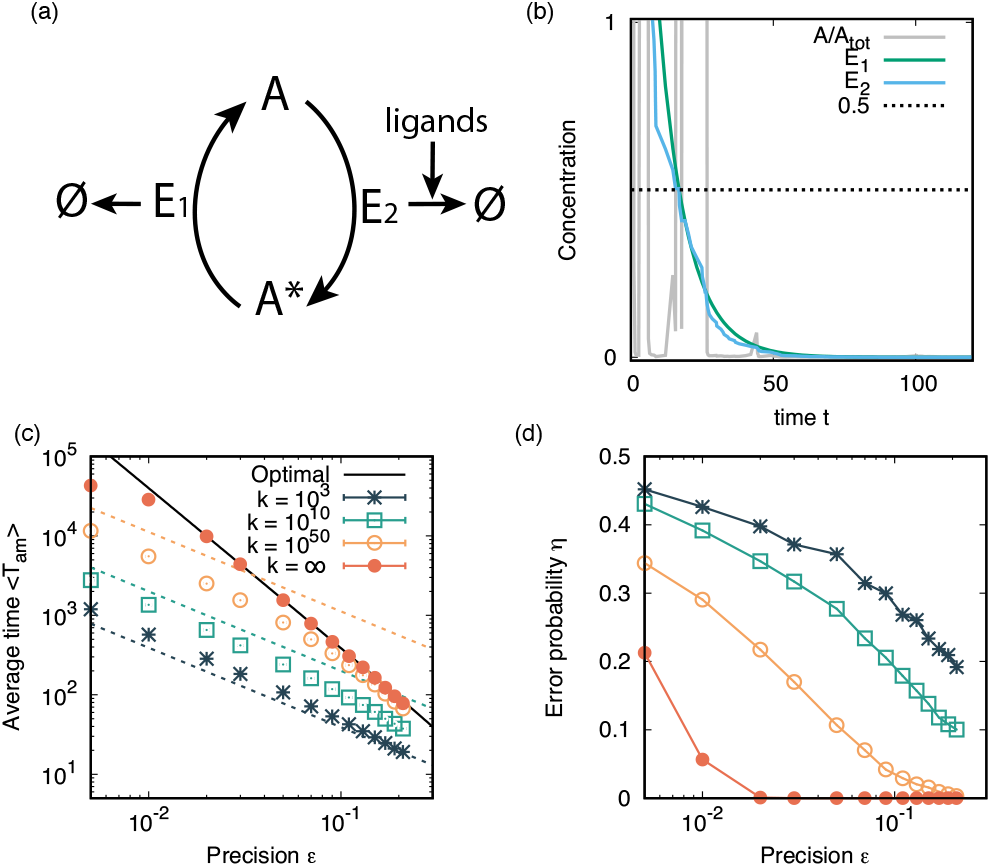
Amendable biochemical model for concentration test. (a) Scheme of the biochemical circuit. (b) A trajectory of the concentrations in the model with *ϵ* = 0.1, *A*_tot_ = 20, ln *E*^(0)^ = 1, *K*_*M*_ = 10^−3^, and *k* = 10^6^, demonstrating the freezing behavior. In this case, *A/A*_tot_ reaches a value < 1/2, representing a correct decision (see Eq. (11)). In all panels, we fixed *τ* = 1/*j* = 1_+_ and varied the precision *ϵ* by varying *j*_−_. (c) Average amendable decision time. Each point represents an average over 10^4^ independent realizations, half with *H* = + and the other half with *H* = −. In all simulations we fixed a final time *t*_*f*_ = 10^5^, *A*_tot_ = 20, *K*_*M*_ = 10 and ln *E* = 1. The optimal theory (black line) is the same as the average amendable time shown in Fig. 6b. Dashed lines represent the characteristic time of freezing *t*^*\**^ estimated by Eq. (12). (d) Error probability. Parameters and meaning of the symbols are the same as for panel (c). Lines joining the points are to guide the eye.

The molecule *A* is a substrate that exists in an unmodified (*A*) or modified (*A*^*\**^) states. These states can be thought of as a dephosphorylated and phosphorylated states, respectively. In this interpretation, *E*_1_ acts as a kinase catalyzing phosphorylation of *A*, while *E*_2_ is a phosphatase catalyzing dephosphorylation. To simplify the model, we assume a quasistatic approximation of the two Michaelis-Menten reactions constituting the switch, so that Eq. (8) reduces to

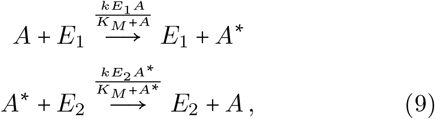

where we also assumed that the two reactions have the same equilibrium constant *K*_*M*_ and production rate *k*. Thanks to this assumption, the GK module switches between a state in which *A* ≫ *A*^*^ and another in which *A* ≫ *A*^*^ when the evidence changes sign, or equivalently when *E*_1_/*E*_2_ = 1 according to Eq. (7). We note that, since *A* + *A*^*^ = *A*_tot_ is constant in time, the kinetics associated with Eq. (9) are governed by a single equation:

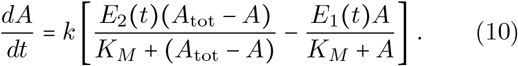

In the regime in which *K*_*M*_ is very small compared to the typical concentrations of *A* and *A*^*\**^, the GK module is characterized by a sharp sigmoidal response. In particular, for large *k*, the quantity 1 − 2*A/A*_tot_ well approximates the sign of the evidence, and can therefore be taken as a proxy for the tentative decision process. We therefore define

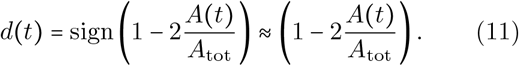

The capacity of producing such sharp response without cooperative binding is the key feature of the GK module. We also note that, since the enzymes *E*_1_ and *E*_2_ decay exponentially, the right hand side of Eq. (10) becomes very small for large time. This means that the dynamics of *A* effectively “freezes” after a certain characteristic time *t*^*\**^, see Fig. 7b. We estimate this characteristic time by noting that, since *K*_*M*_ is small and assuming the difference *E*_2_(*t*) − *E*_1_(*t*) to be on the order of *E*_1_(*t*), the right hand side of Eq. (10) is on the order of *kE*_1_ *t* . In order to accurately track the evidence, this rate should be sufficiently large to induce changes of *A* on the order of *A*_tot_ on the timescale *τ* of individual incoming molecules. We therefore impose *kE*_1_(*t*^*^) ∼ *A*_tot_/*τ*, obtaining

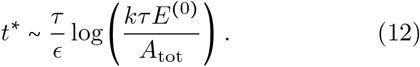

Simulations show that the amendable decision time for *k* → ∞ is in excellent agreement with the optimal theory presented in Section V, see Fig. 7c. Here, the amendable decision time is defined as the last time at which the tentative decision *d t* defined in Eq. (11) switches sign. For finite *k*, the decision process is characterized by a smaller decision time and a higher error rate, see Fig. 7d. This is due to the fact that *A* does not track the evidence sufficiently fast after the time *t*, which is typically much smaller than the optimal decision time. In support of this explanation, the estimate of *t* given in Eq. (12) provides a very good estimate for the amendable decision time for moderately large *k*, see Fig. 7c. Our model demonstrates that biochemical implementation of amendable decisions is feasible, and crucially depends on the capacity of the downstream switch to rapidly track the decaying enzymes. Moreover, this implementation requires a simpler reaction network than for irrevocable decisions.

## VII. AMENDABLE MODEL PREDICTS HUNCHBACK EXPRESSION PROFILE IN DROSOPHILA EMBRYOS

We apply our theory of amendable decisions for concentration tests to the Hunchback-Bicoid system in the fruit fly. Expression of the Hunchback gene represents the first major cell-fate decision made by nuclei in *Drosophila* embryos [27]. This decision is triggered by positional information provided by the transcription factor Bicoid. Specifically, during the nuclear cycles 9 to 13 of the fruit fly development, the transcription factor Bicoid forms an exponentially decaying expression profile along the anterior-posterior axis of the embryo [8, 45– 47]. Hunchback is a gene activated by Bicoid that develops an approximately on-off expression profile depending on whether nuclei are situated in the anterior or posterior half of the embryo, respectively [45, 46, 48, 49], see Fig. 8a. Live imaging of nascent RNA transcription reveals that nuclei in the posterior half sometimes switch off Hunchback expression following a transient activation [50–52] supporting the idea of an amendable decision process.

**FIG. 8.**
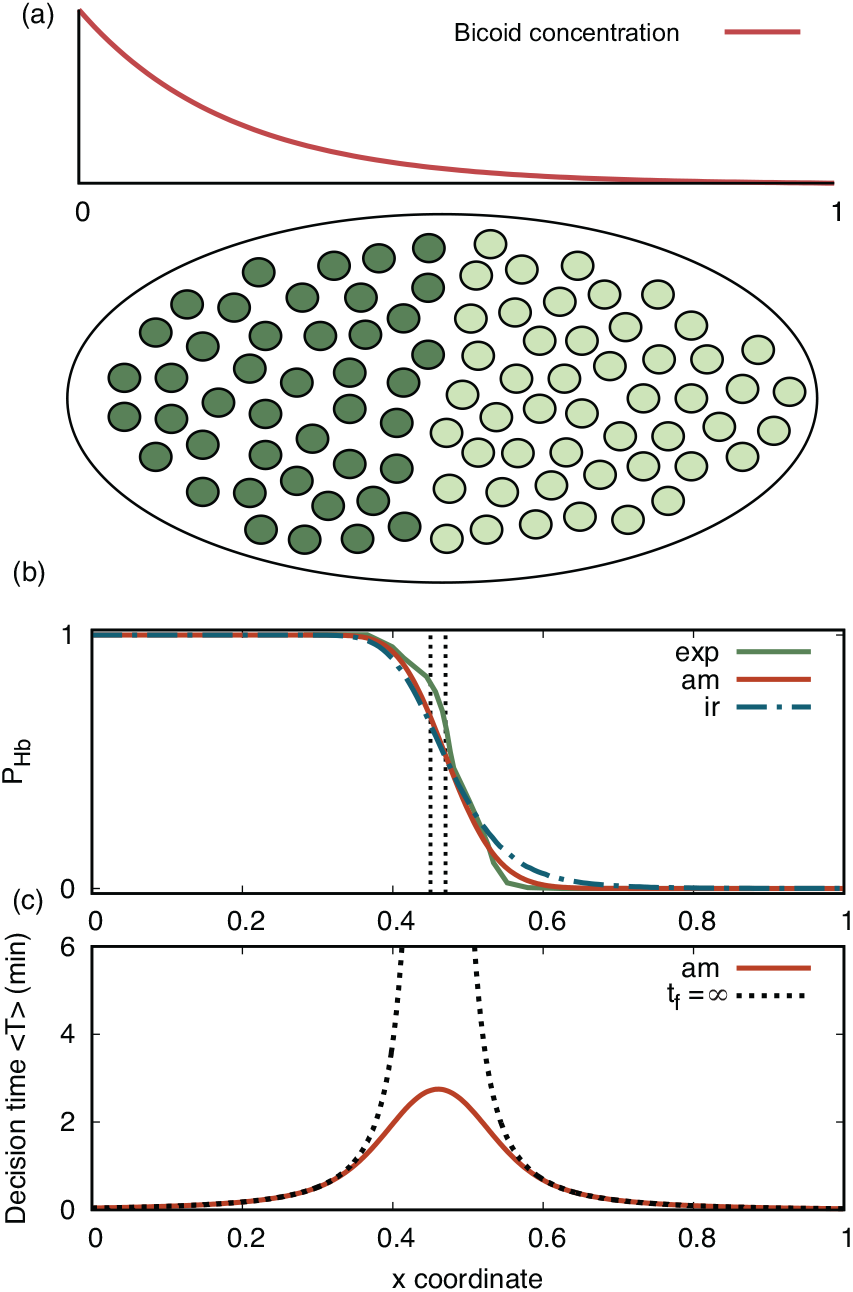
Amendable decisions in the Bicoid-Hunchback system of *Drosophila* embryos. (a) The morphogen Bicoid forms a longitudinal concentration gradient along the embryo. Nuclei sense the Bicoid concentration and decide whether to express Hunchback (anterior half, green circles) or not (posterior half, light circles). (b) Probability *p*_Hb_ to express Hunchback at the end of the 10-th nuclear cycle as a function of the relative coordinate *x* along the anterior-posterior axis. Probabilities of Hunchback expression for amendable decisions (red solid line), compared with experimental data (green solid line) and optimal irrevocable decisions (blue dashed dotted line). Parameters are chosen based on experimental observations (see Appendix L). Experimental data represent the relative number of nuclei that express Hunchback, as identified through fluorescent in situ hybridization (data from Ref. [48]), see also Appendix M. The two dotted vertical lines mark the separation between two neighboring nuclei at the Hunchback interface. Results for 11th, 12th, and 13th nuclear cycle are qualitatively similar, see Fig. 13 in Appendix M. For amendable decisions there is one fitting parameter, namely the Hunchback boundary position *x*_m_, where the expression probability crosses 1/2, while for the irrevocable case we have additionally the two decision thresholds as fitting parameters, see Appendix L. In cases where neither of the thresholds has been reached at the final time, we assumed that cells do not express Hunchback, as in the case in which the negative threshold is reached. (c) Average decision time for amendable decisions based on the optimal evidence process and using the continuous approximation. Results for *t*_f_ = 5.5min (solid line) are compared with the case *t*_f_ = ∞ (dotted line).

To apply our theory to this system, we assume that nuclei are evolutionary adapted to distinguish the concentration differences experienced by neighboring nuclei at the Hunchback boundary position *x*_*m*_, i.e., the point along the embryo axis where Hunchback switches its expression. Based on this assumption, we consider amendable decisions that use the evidence process (5) with the two point processes *N*_+_ and *N*_−_ corresponding with the two Bicoid concentration values left and right of the *x*_m_, respectively (see Appendix L for details). This amendable decision model is able to generate a sharp spatial activation profile of the Hunchback gene in a timescale compatible with that of nuclear cycles, see Fig. 8b. The predicted profile for nuclear Hunchback activation is in excellent quantitative agreement with the experimentally measured one [48] with one fitting parameter (the *x*_m_ coordinate), supporting that the Bicoid-Hunchback system operates in a near-optimal way. The optimal model based on irrevocable decisions leads to a slightly worse fit, but still compatible with experimental data (see Fig. 8b), and requires careful fitting of the decision thresholds (see Appendix L). The decision times predicted by the amendable model are in the order of minutes, and decrease with the distance from the boundary (see Fig. 8c), in qualitative agreement with experiments, see Ref. [51] and Appendix M.

## VIII. AMENDABLE TRANSCRIPTION MODEL

We now study a biophysical model of transcription regulation, to assess whether its dynamics could be thought of as a way to implement reliable amendable decisions. We choose the model architecture and parameters to describe the Bicoid-Hunchback system, although the model can be straightforwardly adapted to other transcriptionally regulated genes.

We consider a promoter that has six binding sites for a transcription factor (Bicoid). The time-dependent number of Bicoid transcription factors bound to the Hunchback promoter constitutes our observation process *X* (*t*), see Fig. 9a. We assume for simplicity that Bicoid binds independently at these six sites, see [53]. Following Ref. [52], we assume that the promoter stochastically switches between ON and OFF states with rates that are biased in favor of the ON state if more than three Bicoid proteins are bound. When the promoter is ON, RNA polymerases (PolII) can bind to the promoter and initiate transcription. We explicitly describe the dynamics of RNA polymerases as a totally asymmetric exclusion process [54]. We identify the evidence process *E* as the sum of the lengths of all Hunchback RNA transcripts bound to PolII minus a constant. This definition is consistent with the fact that the signal measured in fluorescent in situ hybridization, such as those in Ref. [48], is determined by localized transcripts in a compact region, presumably the Hunchback locus. In this interpretation, the subtracted constant represents an experimental detection threshold. The evidence defined in this way tends to increase or decrease in time when the gene is ON or OFF, respectively. It thereby provides nuclei with a mechanism for evidence accumulation. The decision variable *D* is taken as the sign of the evidence at the final time.

**FIG. 9.**
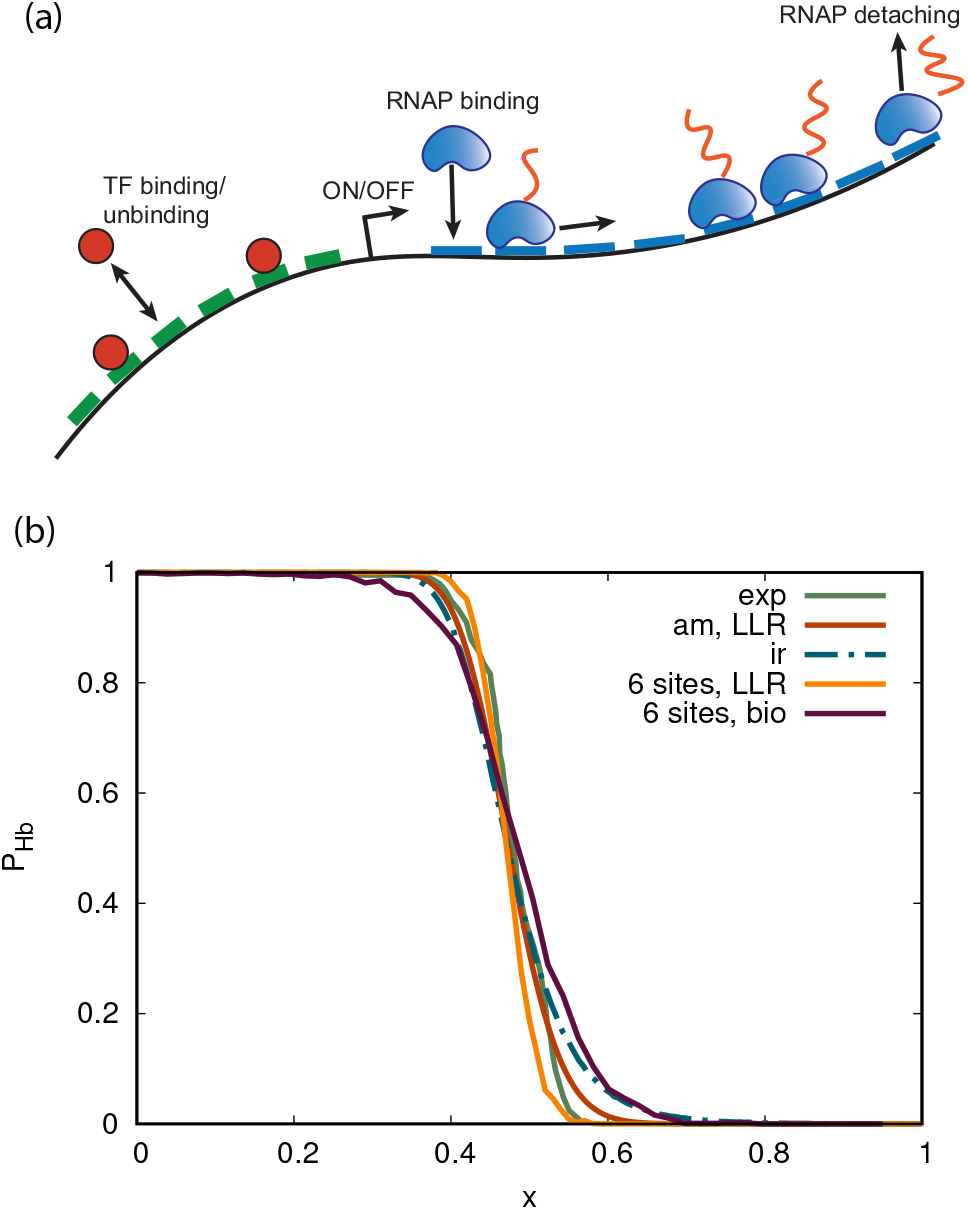
(a) Illustration of the amendable transcription model. (b) Comparison between the probabilities *p*_Hb_ of expressing Hunchback RNA transcripts in the biophysical model and in the corresponding amendable decision model based on the log likelihood ratio (LLR, both averages over 1000 simulation runs). The evidence process is given by Eq. (N5), and in the LLR case the evidence process is governed by Eq. (N7). Parameter choices are presented in Appendix N. For comparison, the data of Fig. 8b is also shown. These include the LLR amendable decision process using the cellular sensing model of Appendix K, the optimal irrevocable decision process, and the experimental data on *p*_Hb_ from Ref. [48].

Figure 9b compares the measured probability of Hunchback activation, shown in Fig. 8b, with two theoretical predictions. The first is the Hunchback activation profile predicted by the biophysical model described in this section. The second is the profile obtained by using as evidence the log-likelihood ratio of the observation process. Unlike the model of Sec. VII, this observation process explicitly accounts for the six Bicoid binding sites, see Appendix N.

The different models and the experimental data are all characterized by similar decision boundaries. The biophysical model predicts a shallower decision boundary than the log-likelihood ratio model, but the difference is small. The gap between the biophysical and normative models can likely be reduced by optimizing free parameters of the biophysical model – an analysis that we leave for a future study.

We remark that the assumptions underlying the model introduced in this section are in line with established models of transcription regulation [52] and dynamics of RNA polymerase [54]. In this respect, our results demonstrate that such dynamics can be interpreted as a way to implement evidence-based amendable decision-making, and lead to a performance that closely approaches the optimal one.

## IX. DISCUSSION

We have shown that amendable decisions are governed by different laws than irrevocable ones. In particular, amendable decisions can be made in a finite average time at zero error probability, thereby offering a substantial advantage over irrevocable decisions. Our results challenge the prevailing paradigm in biology that assumes that decisions, once made, cannot be revoked.

We confirmed the difference between amendable and irrevocable decisions in a visual perception experiment, inspired by classic studies [6, 55]. Our experiment confirmed the possibility of error-free amendable decisions in a finite time, as predicted by the theory. The amendable decision model provides an alternative approach for estimating the evidence acquisition rate. This quantity, usually estimated using irrevocable decision tasks [6], is important in behavioral psychology to measure the impact of factors such as aging [56, 57], aphasia [58], and attention-deficit hyperactivity disorder [59] on cognitive decline. Our study also paves the way for rationalizing amendable decision behavior in more complex cognitive tasks, such as brightness discrimination [60].

Our theory also applies to the microscopic world of cell-fate decisions. For the Bicoid-Hunchback system, we have shown that an amendable decision model that acquires evidence through the likelihood ratio process predicts a decision profile along the embryo in excellent quantitative agreement with experimental data and with only one fitting parameter. Previous studies have observed that the Bicoid-Hunchback system operates close to the optimal physical limit [8], see also [61]. Our theory corroborates this view, and additionally suggests that this limit is reached by an amendable decision process. Biophysical models of the Bicoid-Hunchback model show that the steepness of the irrevocable decision profile can be improved by cooperative binding at the Hunchback promoter [23, 53]. In the amendable case, we have shown that a model of RNA polymerases binding and unbinding provides a simple and biophysically sound mechanism for evidence accumulation, in the form of the total amount of nascent RNA transcripts bound to polymerases. Despite its simplicity, this model predicts realistic Hunchback activation profiles, even in the absence of cooperative binding.

Our theory can shed light on a broader range of cell-fate decisions, with a particularly exciting potential in the study of cell cycle checkpoints. Cells experiencing severe stress or irreparable DNA damage can arrest their cell cycle. Understanding the conditions underlying these decisions is crucial for cancer biology [62]. Decisions to delay or arrest the cell cycle are highly dependent on cellular context and history [63, 64] and must occur in certain time windows. Cells that have arrested their cycle may re-enter it under certain conditions [62, 65, 66], supporting that these decisions are amendable. The theoretical framework developed here can reveal whether these decisions are optimally made.

Broader applications of amendable decisions will require going beyond the paradigm of binary hypotheses. Examples are change-point detection problems, of potential relevance for accurate timing of developmental cell-fate decisions [67], and multiple hypotheses sequential testing [13] for modeling cell differentiation in higher organisms.

The superior performance of amendable decisions over irrevocable ones raises the question of why certain decision processes in biology are hardly reversed. One way of rationalizing the distinction between amendable and irrevocable decisions is to compare the cost of making an error with the cost of correcting it. From this perspective, amendable and irrevocable decisions represent two extremes of a continuous spectrum: at one end, the cost of amending a decision is negligible relative to the cost of making an error; at the other end, the cost of amendment outweighs the consequences of an error. During early development, errors pose a particularly severe risk due to error amplification [27], suggesting that amendable decisions may serve as a natural control mechanism.

## ACKNOWLEDGMENTS

IN conducted part of this research while visiting the Okinawa Institute of Science and Technology (OIST) through the Theoretical Sciences Visiting Program (TSVP). We thank A. Sassi for assistance with the visual perception experiment. We thank E. Kanemoto and G. Tripp for feedback on the experimental protocol, H. Sprekeler for comments on the drift-diffusion model, and S. Bo, N. Luscombe, and G. Tkacik for feedback on a preliminary version of the manuscript. The final version of the manuscript was proofread for typos with the help of Claude Opus 5.5.

## Appendix A: Drift-diffusion model: irrevocable decisions

We here present general mathematical results for irrevocable decisions based on a drift-diffusion evidence process. We assume that the evidence is a linear function of the observation process *X*, and thus, for a given hypothesis, obeys the dynamics

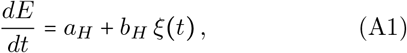

with an initial condition *E* (0) = *E*_0_, and where *a*_*H*_ and *b*_*H*_ are constants that depend on which of the two hypotheses holds (*H* ∈ {+, −}). Equation (A1) is more general than Eq. (2), as the latter assumes that the average evidence rate is odd under switching hypothesis. This assumption always holds if the evidence process is chosen as the log-likelihood process (as we shall prove in Appendix C). The assumption can also be justified by a symmetry argument in cases when the two hypothesis are related by a change of sign, *µ*_+_ = −*µ*_−_, as in our visual perception experiment.

We still consider for simplicity symmetric thresholds, *l*_+_ =*l*_−_ = *l*; we notice that for drift-diffusion processes first-passage problems with asymmetric thresholds can be mapped on first-passage problems with symmetric threshold by changing the initial value *E*_0_. The probabilities of reaching the and thresholds, given the ± hypotheses, are given by

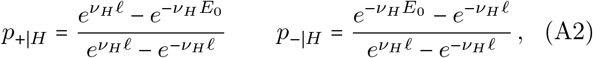

respectively, where 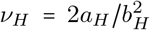. We can derive this result by using that *M*_*H*_(*t*) = exp(−*v*_*H*_*E*(*t*)) is a martingale and applying Doob’s optional stopping theorem [68–70]. For *E* 0 0, Eq. (A2) reduces to

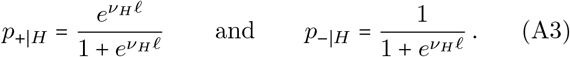

We compute the distribution *p*_ir_(*t*) of first-passage times from the backward Kolmogorov equation [71], obtaining

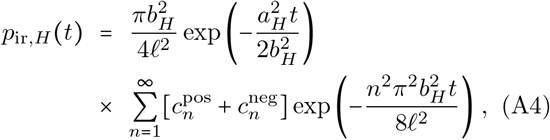

where

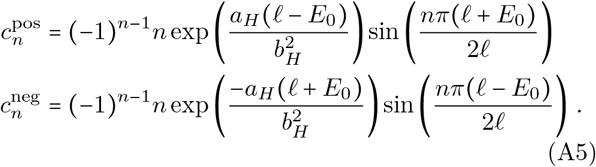

The coefficients 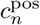 and 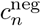 correspond to contributions originating from the positive and negative threshold, respectively. Also in this case, the distributions of decision times have a simpler expression for *E*_0_ = 0:

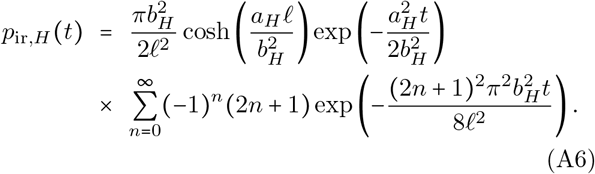

We express the average decision time by means of Wald’s equation [69, 72, 73], obtaining

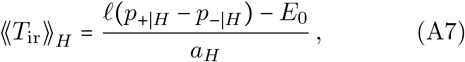

where we have introduced the notation ⟪…⟫_*H*_ for an average over the observation process without averaging over the hypotheses.

The error probability *η* is the sum of the probabilities of false positives and false negatives:

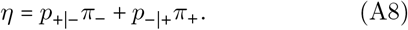

where we introduced the prior probabilities *π*_+_ and *π*_−_ of the plus and minus hypotheses, respectively.

The threshold *l* is often chosen according to the desired value of the error probability. For example, in the case of a symmetric evidence process (Eq. (2), such that *a*_±_ = ±*γ* and *b*_±_ = 1) and uniform prior (*π*_+_ = *π*_−_ = 1/2), Eq. (A3) leads to

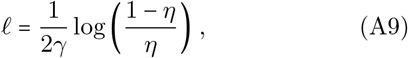

so that

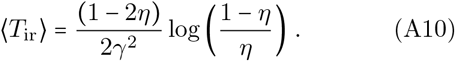

Equation (A10) leads to the speed-accuracy tradeoff relation ⟨*T*_ir_⟩ ∼ − ln (*η*), valid for small *η* [1].

Results for the optimal irrevocable decision strategy, where the evidence process is proportional to the logarithm of the likelihood ratio, are recovered by setting 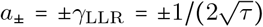 and *b*_±_ = 1 in the equations of this section.

## Appendix B: Drift-diffusion model: amendable decisions

In the amendable case, the decision time is defined as the last time that the evidence *E* has changed sign. We directly consider the general case in which the evidence is governed by Eq. (A1) and decisions are made within a finite time window [0, *t*_f_]. We denote by *S*_*H*_ (*E*_0_, *t*) the probability that the process, starting from *E* = *E*_0_ at time 0, does not reach *E* = 0 by time *t*, depending on the hypothesis *H* ∈ {+, −}. This probability is expressed by

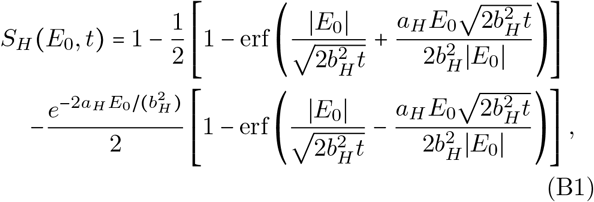

for *E*_0_ ≠ 0 and *S*_*H*_ (0, *t*) = 0. Here, erf(*x*) = 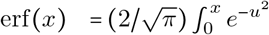 *du* denotes the error function, see, e.g.,Section (3.2.2) in Ref. [39].

Our aim is to compute the probability *s*_±(_*t*) that a zero crossing at time *t* is the last one. To this aim, we consider *S*_±_ (*ϵ*_*E*_, *t*) for small |*ϵ*_*E*_|. Starting from *E* = 0 a fluctuation leads to *E* = *ϵ*_*E*_ or *E* = −*ϵ*_*E*_ with equal probability due to the properties of Brownian motion. Thus, the escape probability per unit evidence takes the form

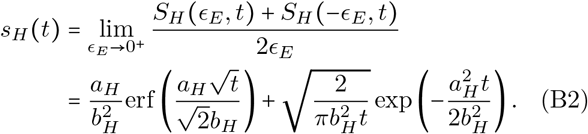

The finite-time decision time distribution is proportional to the probability of being in state *E* = 0 at time *t*, times the probability that the process will not return to the state *E* = 0 by time *t*_f_ :

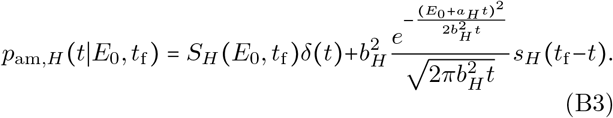

The first term on the right-hand side of Eq. (B3) is the contribution from the trajectories that never cross zero. The second term is proportional to the probability of being in *E* = 0 at a given time *t*, times the probability of not returning to zero again before the final time. The prefactor 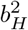 in the second term in the right hand side can be either justified by a dimensional argument, or by numerically verifying that only with this choice the distribution in Eq. (B3) is correctly normalized to one.

At finite times, amendable decisions entail a finite error probability. If the decision process is reliable, then this probability is simply given by the probability that, at the final time, the sign of the evidence *E* is different than the sign of the bias *a*. Since the distribution of *E* is Gaussian at any time, we find

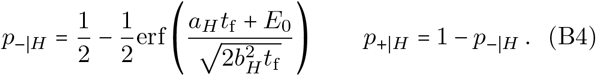

In the limit *t*_f_, we obtain a vanishing error probability and a distribution of decision times given by

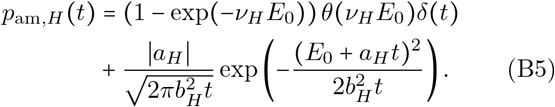

We recall that *v*_*H*_ 2*a*_*H*_ *b*^2^ . The corresponding average decision time is expressed by

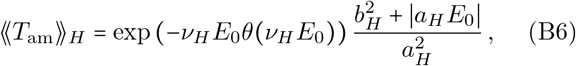

which reduces to 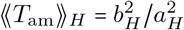 for *E*_0_ = 0.

We note that for *a*_*H*_ = 0 and *E*_0_ = 0, Eq. (B3) reduces to 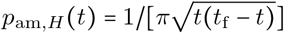, which is the second arcsine law for Brownian motion [74].

## Appendix C: Log-likelihood as an evidence process

In this appendix, we consider an evidence process that is a generic additive functional of a drift-diffusion observation process (Eq. (1)) and we derive the condition for the evidence process to be proportional to the log likelihood.

Assuming that the evidence process is an additive functional of the observation process, we obtain

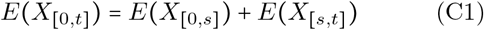

for all *s* ∈ [0, *t*]. Here, *X*_[0,*t*]_ denotes the trajectory of *X* in the time interval [0, *t*]. The assumption of time-additivity can be justified by the fact that *X* is a Markovian drift-diffusion process. Using that *X t* is time-homogeneous, this implies that *E* evolves in time as

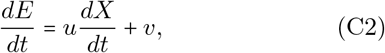

where *u* and *v* are two constants. Substituting Eq. (1) into Eq. (C2), we find

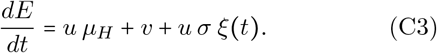

As Eq. (C3) is invariant under a simultaneous rescaling of *u, v*, and the units of *E*, we set without loss of generality the noise amplitude to one by imposing *u* = 1/*σ*, obtaining

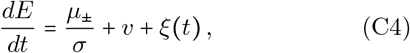

which is a special case of Eq. (A1). In keeping with Eq. (A1), we therefore have

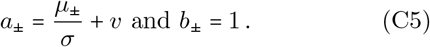

Given the observation process, the parameter *v* is the only remaining degree of freedom. This parameter does not affect the difference of the two drifts, *a*_+_ − *a*_−_ = (*µ*_+_ − *µ*_−_)/*σ* = 1/ *τ* = 2*γ*_LLR_.

We now show that Eq. (2) with *γ γ*_LLR_ is satisfied by an evidence process

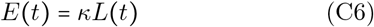

proportional to the log-likelihood ratio process

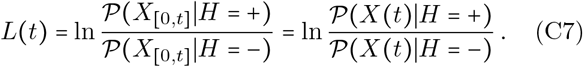

Here, *P*(*X*_[0,*t*]_|+) is the probability distribution of observing the full trajectory *X*_[0,*t*]_ conditioned on *H* = +, *P*(*X*_[0,*t*]_|−) is the corresponding distribution for *H* = −, *κ* is a proportionality constant, and the last equality follows from the fact that *X*(*t*) is a drift-diffusion process. Using that the probability distribution of *X* is a Gaussian with mean *µ*_±_*t* (depending on the hypothesis) and variance *σ*^2^*t*, we obtain

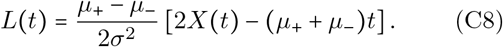

Taking a time derivative and using Eq. (1) we find

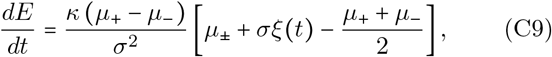

where *κ* = *σ/*(*µ*_+_ − *µ*_−_). Substituting *κ* in the above evolution equation we obtain, after some algebra,

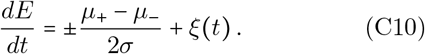

Equation (C10) corresponds to Eq. (C4) for

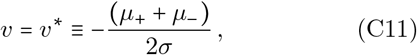

and also to Eq. (2) for *γ γ*_LLR_, as we were meant to show.

We note that the choice *v v* corresponds to the symmetric splitting

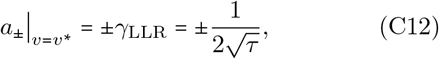

i.e., the log-likelihood ratio process is the unique member of the family of evidence processes defined by Eq. (C4) that treats the two hypotheses symmetrically.

## Appendix D: Optimality for irrevocable decisions

For completeness, we briefly summarize the theory for optimality of irrevocable decisions in sequential hypothesis testing [12, 13]. For irrevocable decisions, we say that the evidence process *E* is optimal if it minimizes the average decision time ⟨*T*_ir_⟩ = *π*_+_ ⟪*T*_ir_⟫_+_ + *π*_−_ ⟪*T*_ir_⟫_−_ over the set of all evidence processes *E* that have prescribed maximal probabilities *α* and *β* of false positives and negatives, respectively:

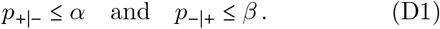

For drift-diffusion observation processes, the optimal evidence process is proportional to log-likelihood ratio process *L*(*t*) (see Eq. (C7)), corresponding with *v* = *v*^*^ in (C4). Specifically, Wald and Wolfowitz have shown in Ref. [40] that, for discrete-time observation processes consisting of independent and identically distributed random variables, *L*(*t*) is optimal if the thresholds are set so that

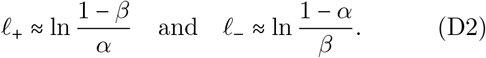

As noted in Ref. [73], the argument of Ref. [40] does not rely much on the discrete time nature of the process, and thus it was conjectured that also in continuous time *L(t)* is optimal. In this case, the relations in Eq. (D2) are equalities. The optimality of the *L(t)* in continuous-time drift-diffusion processes was formally proven by Shiryaev in Section 4.2 of his book [75], and later generalized to continuous-time processes with stationary and independent increments in Ref. [76].

For a general observation process *X*(*t*), the optimal evidence process *E*(*t*) for irrevocable decisions is not known. In the asymptotic limit of vanishing error probabilities, and thus for large *l*_+_ and *l*_−_, one can consider the easier optimization problem of minimizing the leading behavior of ⟨*T*_ir_⟩ as a function of *l*_+_ and *l*_−_. It was demonstrated that *L*(*t*) is asymptotically optimal given a suitable convergence condition on the process *L*(*t*)/*t*, see Theorem 3.4.2 on page 167 of Ref. [13]. Specifically, it is required that *L*(*t*)*/t* converges *r*-quickly with *r* ≥ 1 to a deterministic limit [77]. This result implies that *L*(*t*) well approximates the optimal process in the limit of small error probability.

## Appendix E: Optimality for amendable decisions

For amendable decisions in an infinite time window, *t*_f_ = ∞, we say that the evidence process *E* is optimal if it minimizes the mean decision time

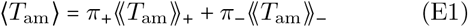

over the set of all evidence processes *E* for which the probability of error vanishes, *η* = 0. We here find the optimal evidence process under this definition.

As a baseline, we first compute the average decision time for the log likelihood evidence process under a uniform prior. Substituting *E*_0_ = 0, *b*_*H*_ = 1 and ∣*a*_*H*_| = (*µ*_+_ − *µ*_−_)/(2*σ*) (see Eq. (C10)) into Eq. (B6) we immediately obtain 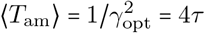, as stated in Section II.

We now formally derive the optimal amendable evidence process, and compare with the baseline given by the log likelihood process. We perform the calculation for *E*_0_ ≥ 0, so that the observer initially prefers the + hypothesis, and then use the symmetry

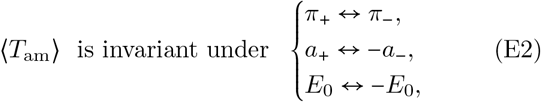

to extend the result to negative values of *E*_0_.

Using Eq. (B6), we find that the mean amendable decision time, associated with a general evidence process that evolves according to Eq. (C4), is given by

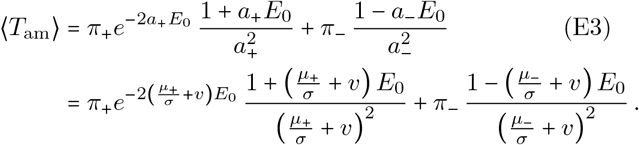

Our aim is to minimize *T*_am_ with respect to *v* and *E*_0_, subject to the constraint that the error probability vanishes, which is equivalent to assuming

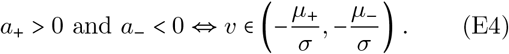

We note that the log-likelihood ratio process corresponds to the midpoint of the admissible interval for *v*, see Eq. (C11). Mathematical expressions in the following are provided in terms of *a*_+_ and *a*_−_ rather than *µ*_+_, *µ*_−_, and *v*, as the former expressions are more compact. We also note that the mean decision time diverges when *v* tends to the boundaries of the admissible interval defined in Eq. (E4). This implies that the optimal value of *v* must necessarily be in the interior of the domain.

Taking the derivative of Eq. (E3) with respect to *E*_0_ we find

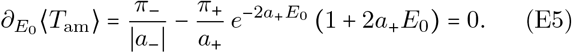

Solving Eq. (E5) towards *E*_0_, we find the solution for positive values of *E*_0_, and using the symmetry (Eq. (E2)) we extend this result then to negative values. This yields the optimal value of *E*_0_ given by

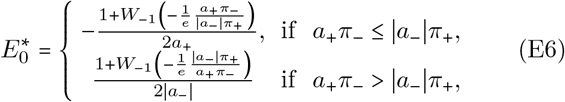

where *W*_−1_ is the lower real branch of the Lambert-W function, solving *W*_−1_(*x*)*e*^*W*−1(*x*)^ = *x* with *W*_−1_(*x*) ≤ −1 for *x* ∈ [−1/*e*, 0)_*_.

To obtain *v*^*\**^, the optimal value of *v*, we impose a vanishing derivative of ⟨*T*_am_⟩ with respect to *v*. Using Eq. (E3) we find

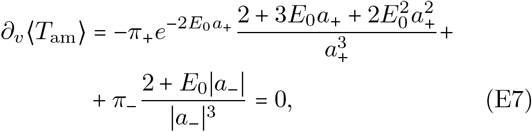

for *E*_0_ ≥ 0, and

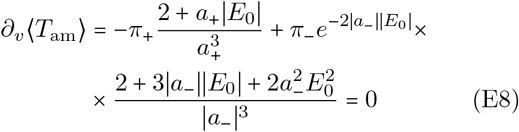

for *E*_0_ < 0. We are thus left to solve a relatively complicated set of nonlinear equations, given by Eqs. (E6) and Eq. (E7).

Numerical solution shows that the minimum average decision time is attained with a value of *v* different from *v*^*^, see Fig. 3. This means that, in general, the optimal evidence process for amendable decisions is not the log likelihood process.

We now study the special case of a uniform prior *π*_+_ = *π*_−_ = 1/2. In this case, Eqs. (E5) and (E7) admit three stationary points, two of which are minima and one a saddle, see Fig. 3. The saddle point corresponds to the log-likelihood ratio process. Indeed, the choice of *v* = *v*^*^ (see Eq. (C11)) together with *E*_0_ = 0 simultaneously solves Eq. (E5) and Eq. (E7), as *µ*_−_ /*σ* + *v* = −(*µ*_+_ /*σ* + *v*) and *W*_−1_(−1/*e*) =−1. Evaluating the second derivatives of Eq. (E3) at (*v*, 0) we find

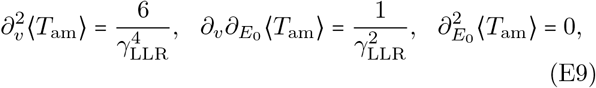

so that the determinant of the Hessian equals 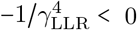, and thus surprisingly this is a saddle point. To find the location of the minima, we consider *E*_0_ ≥ 0. Equations (E5) and (E7) read

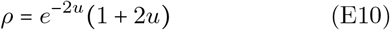

and

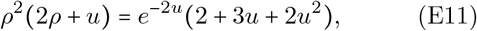

respectively, with *ρ* = *a*_+_/∣*a*_−_∣ and *u* = *a*_+_*E*_0_. Eliminating *ρ* from Eq. (E11) and setting *y* = *e*^−2*u*^, we find the quadratic equation

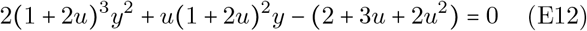

and its positive root is given by

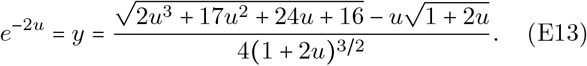

Equation (E13) is a self-consistent equation in *u*, with solutions *u =* 0 and *u* ≈ 0.2272. From Eq. (E3) we obtain

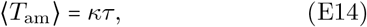

where the prefactor is

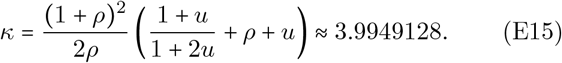

We recall that the log-likelihood ratio test provides an average amendable decision time equal to 4*τ*, which is thus very close to the optimum.

## Appendix F: Minimax optimality for amendable decisions

In this appendix we consider minimax optimality, an alternative definition of an optimal decision as the process that minimizes

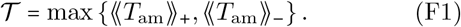

We shall prove that

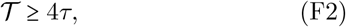

and one readily verifies that the equality is attained for the log-likelihood ratio process with (*v, E*_0_) = (*v*^*^, 0), for which ⟪*T*_am_⟫_+_ = ⟪*T*_am_⟫_−_ = 4*τ* . Since *T* does not involve the prior, this conclusion holds for arbitrary *π*_±_.

In what follows we prove the inequality *T* ≥ 4*τ* . Thanks to the mirror symmetry expressed in Eq. (E2), we set *E*_0_ ≥ 0 without loss of generality. Furthermore, we assume the conditions given in Eq. (E4) so the error probability equals zero, and we set *b*_*H*_ = 1. Since 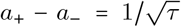 is fixed, we parametrize the admissible evidence processes by a single dimensionless number 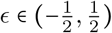,

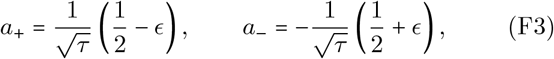

with *ϵ* = 0 corresponding to *v* = *v*^*^, together with *u* ≡ *a*_+_*E*_0_ ≥ 0. Equation (B6) then reads

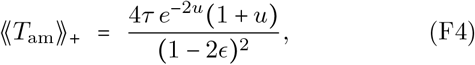

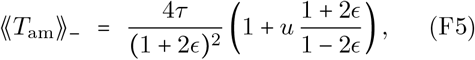

where we used ∣*a*_−_∣ *E*_0_ = *u* (1 + 2*ϵ*) */* (1 − 2*ϵ*).

Since *T* is a maximum, it suffices to show that at least one of the two expressions, Eq. (F4) or Eq. (F5), is larger than or equal to 4*τ* . We determine the values of *u* for which this condition holds. For Eq. (F5) this is immediate:

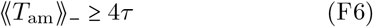

whenever 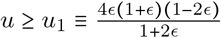 . For Eq. (F4) we use the inequality *e*^−2*u*^(1 + *u*) ≥ 1 − *u*, valid for *u* ≥ 0 because *h*(*u*) = *e*^−2*u*^ (1 + *u*) − 1 + *u* satisfies *h*(0) = *h*′ (0) = 0 and *h*^′′^(*u*) = 4*ue*^−2*u*^ ≥ 0, the inequality being strict for *u* > 0. We obtain

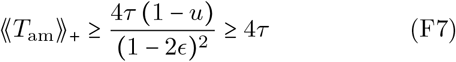

whenever *u* ≤ *u*_2_ ≡ 1 – (1 − 2*ϵ*) ^2^ = 4*ϵ* (1 − *ϵ*). The two conditions cover all admissible *u*, because

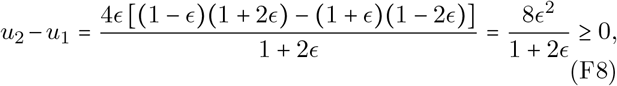

so that *u*_1_ ≤ *u*_2_. Hence *T* ≥ 4*τ* everywhere in the admissible region, completing the proof.

## Appendix G: Visual perception experiment: protocol

Eight volunteering participants were selected among students and researchers at OIST. Each participant repeatedly played a game in which a dot randomly moves on a computer screen, with the goal to assess whether the motion was biased towards the left or right direction and make decisions using the computer keyboard.

The game had an irrevocable and amendable version. In the irrevocable version, the game ended once a decision was taken, while in the amendable version participants could alter their decision an arbitrary number of times, up to a time limit of 10s. For both types of games, participants were asked to decide rapidly while avoiding mistakes.

Each participant played eight groups of 30 games each. Four out of eight groups consisted of irrevocable games and the remaining four of amendable games. Irrevocable and amendable groups of games were played in an alternate fashion. Half of the participants started with an irrevocable group and half with an amendable group, to remove potential biases. The four groups for each game involved an increasing level of difficulty. Specifically, we express as (1/2 + *q*) and (1/2 – *q*) the probabilities of moving in the direction of the bias or in the opposite direction, respectively, during a timestep Δ*t =* (1/60) s of the game. The bias parameter *q* ranged in the set of values *q* ∈ {0.15, 0.11, 0.07, 0.03} for the four games. The corresponding parameters of the observation process are *µ*_±_ = ±2*q*/Δ*t* and *σ*^2^ = (1 − 4*q*^2^)/Δ*t*. The dot also performed unbiased Brownian motion in the vertical direction.

Each of the 8 participants performed the experiment in a single session. Participants were asked to read an instruction sheet outlining the description of the amendable and irrevocable games. The instruction sheet also included a consent agreement. Participants were then given an opportunity to ask questions to clarify possible doubts. All participants provided informed consent.

Before the actual experiment, participants had the opportunity to practice both irrevocable and amendable games. They first could practice these games in a simplified version, in which the initial position of the dot was marked with a vertical dashed line. They then could practice the normal version of the game, without the reference line. For all the practice games, the difficulty was set to *q =* 0.07. Each version of the practice game could be played for as long as the participants felt comfortable with. Both in the practice and actual experiment, the game provided summary information at the end of each round, such as whether the decision was correct or incorrect, the decision time, and the average decision time. During the actual experiment, participants were encouraged to take breaks of a few minutes between the 4 × 2 = 8 groups of games. The entire session lasted about one hour per participant, including the introduction and practice. Participants were anonymized before the data analysis. The study protocol was approved by the OIST human subject research committee (reference number HSR-2024-016-2).

## Appendix H: Visual perception experiment: data analysis

In the optimal (normative) model, we obtain the evidence rate from the drift and diffusion coefficients characterizing the dot motion: 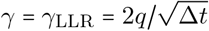. This parameter fully specifies the optimal amendable model. For the optimal irrevocable model, we estimate the decision threshold *l* from the average empirical error probability through Eq. (A9).

In the descriptive model, the evidence rate is estimated from the experimental data. This estimate is different in the amendable and irrevocable cases. For amendable decisions, we combine Eq. (B6) and Eq. (2), obtaining 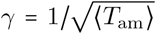. This formula permits to simply estimate the evidence rate from the average decision time. For the hardest game (*q* = 0.03), we took finite-time effects into account to estimate the drift. This can be done either from the mean decision time (obtained by numerically integrating Eq. (B3)) or from the error probability, Eq. (B4). To be conservative, we took the smallest of these two values for each point. In the irrevocable case, we estimate the evidence rate by solving the expression of ⟨*T*_ir_⟩ and *η* towards *γ* and *l*. This procedure leads to 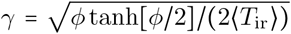 and 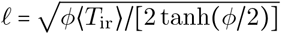, where *ϕ* = log[*η*/(1 − *η*)].

## Appendix I: Visual perception experiment: individual speed-error performance

In this Appendix, we perform a more detailed analysis of the visual perception experiment at the individual level. We compare the average amendable decision time for each participant and each game difficulty with the optimal prediction, see Fig. 10. We consider two different types of optimal predictions. The first uses as evidence the logarithm of the likelihood ratio process, and is evaluated on the same observation process as seen by the participants. For convenience, in this section we describe this observation process as a discrete-time random walk, rather than a continuous time drift-diffusion process. This correspond to the fact that the dot moves in fixed time steps Δ*t* = 1/60s one pixel to the left or right. Correspondingly, the normative evidence process is the likelihood ratio process of a discrete random walker. The second optimal prediction uses the theoretical expressions for the mean amendable decision time and the error probability of a drift-diffusion process.

**FIG. 10.**
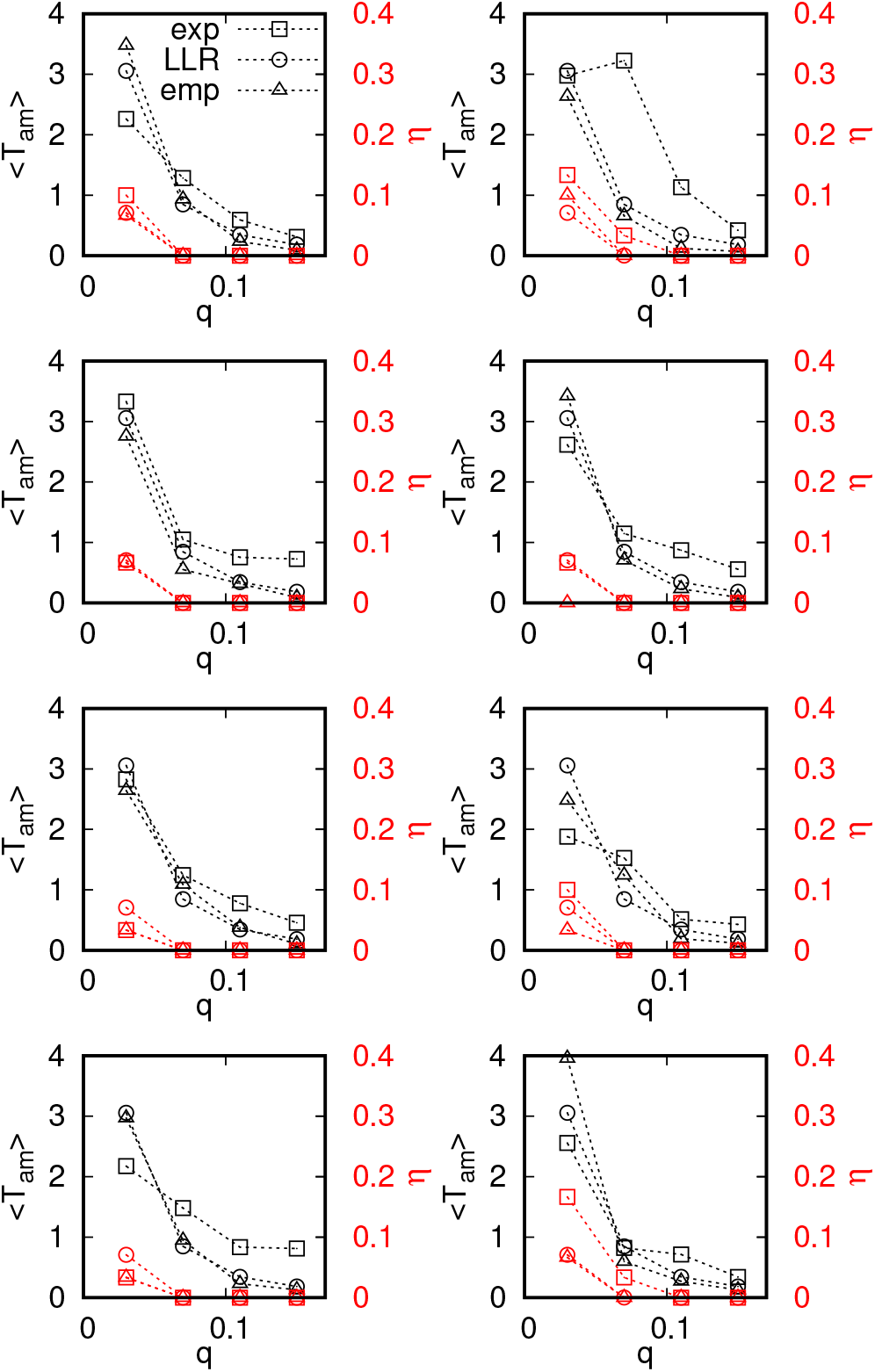
Mean decision times (black) and error probabilities (red) in the amendable games for each of the participants as a function of the game difficulty parameter *q*. Plots compare the experimental data averaged over 30 games (squares) with the mean decision times of an observer that sees the same data as the participants but uses the log-likelihood ratio strategy (triangles) and the mean decision time based on the log-likelihood ratio for infinite sample sizes (circles). Each panel represents one participant (left: participants 1 to 4 and right: participants 5 to 8 top to bottom).

Specifically, the horizontal dynamics of the dot is governed by the dynamics

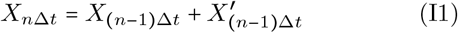

where 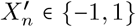 are independent and identically distributed random variables with 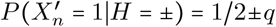, and 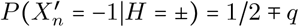, and where *q* is the bias parameter. The time step is Δ*t* = 1/60s and the maximum game duration is *t*_f_ = 10*s*. The associated optimal strategy is based on the evidence process

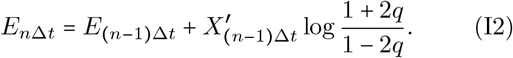

We compared the empirical mean, amendable decision times, and the probability of error for each of the participants as a function of the game difficulty parameter *q*, see Fig. 10. The empirical values are averaged over the 30 games played by each participant for each difficulty level. These empirical values are compared with the optimal mean decision time and error obtained with the optimal decision strategy (using the likelihood ratio as the evidence) for either the drift diffusion process or for the same sample as seen by the participants, see Fig. 10.

Specifically, the amendable decision time for one sample is given by

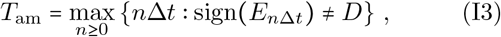

and an error is made if *D* sign (*E*_600Δ*t*_) ≠ *H*. The black and red markers represent, respectively, the mean decision times and the error for the average log-likelihood ratio strategy in continuous time, as considered in Section IV. The error is expressed by

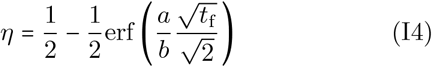

and the mean decision time is the average of the distribution given in Eq. (B3), both with *t*_f_ = 10 and 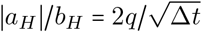.

We recall that participants started with the easiest game (*q* = 0.15), and alternated amendable with irrevocable decision games. Participants 1, 3, 5, and 8 initiated the experiment with an amendable decision game, while participants 2, 4, 6, and 7 initiated the experiment with an irrevocable game.

The analysis presented in this appendix permits to appreciate the individual differences among participants (which are qualitatively consistent with similar studies in the literature) and additionally shows that the optimal average strategy in the continuous limit is broadly consistent with the more detailed optimal strategy for individual samples in discrete time.

## Appendix J: Visual perception experiment: code

The code used for the visual experiment is provided on GitHub [78]). The code is written in Python and requires installation of the library Pygame (see www.pygame.org).

The practice rounds can be launched using the command python3 practice.py. This executable permits to select four practice games: irrevocable with aid, irrevocable without aid, amendable with aid, amendable without aid. Each of these games can be played for an arbitrary number of rounds. Pressing q exits the game to the selection screen.

The actual experiment session is launched with either python3 movingdDotGameA.py or python3 movingdDotGameB.py. The only difference between the two is the order of the games: movingdDotGameA.py starts with an irrevocable game, whereas movingdDotGameB.py starts with an amendable game. As discussed in Appendix G, half of the volunteers played movingdDotGameA.py while the other half played movingdDotGameB.py. In both cases, games alternate while the difficulty level increases as described in Appendix G.

## Appendix K: Amendable decision times for the cellular sensing model

Our starting point is the equation governing the evidence

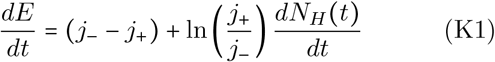

for *j*_+_ > *j*_−_ and *H* ∈ {+, −}, see [28], and the initial condition *E*_0_ = 0. Expressing *j*_−_ and *j*_+_ in terms of the precision parameter *ϵ* = (*c*_+_ − *c*_−_)/*c*_+_ = (*j*_+_ − *j*_−_)/*j*_+_ and the characteristic time scale *τ* = 1/*j*_+_ directly leads to

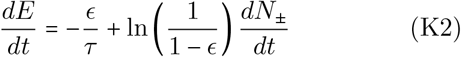

which is Eq. (5). We assume without loss of generality that *ϵ ∈* 0, 1 .

Trajectories of the optimal evidence process are characterized by a deterministic linear decrease with slope −*ϵ*/*τ* that is occasionally interrupted by stochastic upward jumps of size ln(1/(1 − *ϵ*)), corresponding to increments in the counting process *N*_±_(*t*), see Fig. 11. Physically, these jumps represent events in which one particle is absorbed by the observer. The fact that jumps are always upwards implies that the statistics of decision times is qualitatively different for the two hypotheses.

**FIG. 11.**
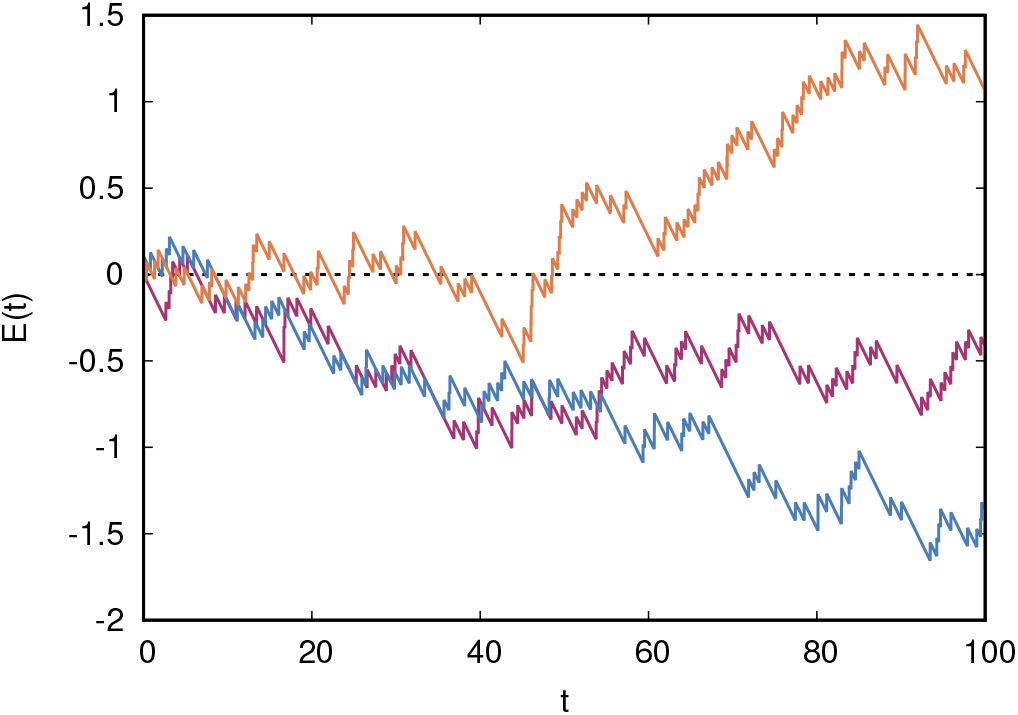
Trajectories of the evidence process defined in Eq. (5) for *ϵ* = 0.1, *τ =* 1, and *H* = +. The negative values of *E*(*t*) are transient and the evidence process drifts eventually upwards.

We study the amendable decision time distribution for *t*_*f*_ = ∞, so that the error probability vanishes. The decision time is the last time when the sign of the evidence was inconsistent with the hypothesis, i.e., sign (*E*(*t*)) = −*H*

We find that the distribution of decision times for the positive hypothesis is expressed by

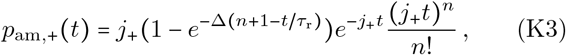

where Δ = ln(*j*_+_/*j*_−_) = ln(1/[1 − *ϵ*]) is the magnitude of the upward jumps, *τ*_r_ = (*τ* /*ϵ*)Δ, and *t* ∈ [*nτ*_r_, (*n* + 1)*τ*_r_]. For the negative hypothesis, we obtain

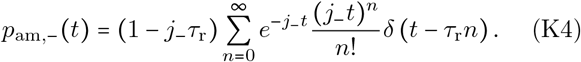

Before proving Eq. (K3) and Eq. (K4), we explore some of their consequences. For the positive hypothesis, we could not find a simple expression for the average decision time. Nevertheless, this time can be obtained from numerically integrating Eq. (K3). The mean time for Eq. (K4) is readily found by summing the series:

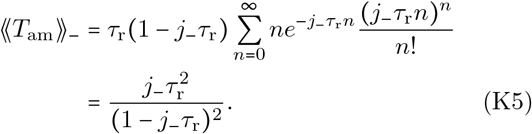

Alternatively, we can use a diffusion approximation of the counting process:

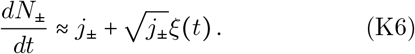

Using Eq. (B6), we then find

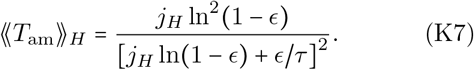

We note that, for the negative hypothesis, the continuous approximation of Eq. (K6) exactly matches the solution given in Eq. (K5). In contrast, for the positive hypothesis, Eq. (K6) only approximates the numerically evaluated mean decision time. Finally, expressing Eqs. (K5) and (K7) in terms of *ϵ* and *τ* we find that ⟪*T*_am_⟫ _*H*_ ∼ *τϵ*^−2^ for small *ϵ*, independently of the hypothesis *H*.

To determine the mean irrevocable decision times ⟪ *T*_irr_⟫ _*H*_ for the two hypotheses, we use again Wald’s equation [70, 72, 73]

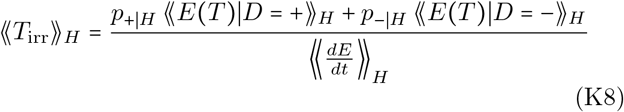

where

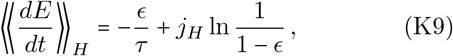

and where we set

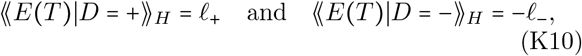

neglecting overshoots. We have verified with simulations that neglecting overshoots yields a very good approximation.

Averaging Eqs. (K7) and (K8) over the two hypotheses and taking the ratio of the two times, we find that this ratio is given by

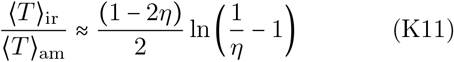

for small values of *ϵ*. From Eq. (K11) we find that amendable decisions are faster than irrevocable ones for *η* ⪅ 0.083, as in the drift-diffusion case (see Fig. 2a) and independently of the precision *ϵ*.

**Proof for** *H* = +. We proceed with the derivation of Eq. (K3). The probability of *T*_am_ ∈ [*t, t* + *dt*] can be expressed by requiring that *E*(*t*) ∈_′_ [−Δ, 0], that after the jump *E* remains positive for all *t* > *t*+ *dt*, and that a particle arrives in the interval *t, t dt* . These requirements lead to

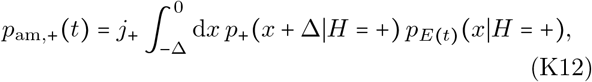

where *p*_+_ (*x* + Δ | *H* = +)is the probability that *E* does not fall below zero given that its initial value is *x* + Δ.

The probability density of *E* at time *t* is fully determined by the number of jumps made by the process up to the time *t* through the formula

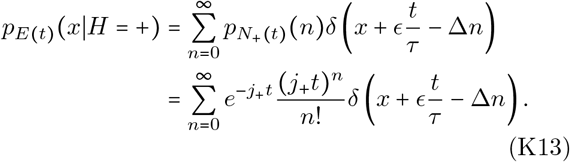

The survival probability *p*_+_(*x* + Δ∣*H* = +) can be determined from the facts that the process exp(−*E*(*t*)) is a martingale [69] and that *E*(*t*) is a continuously decreasing function intercepted by upward jumps. Indeed, consider the first-passage problem

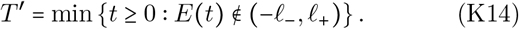

Applying Doob’s optional stopping theorem to the martingale exp(−*E*(*t*)) [68, 69] we find 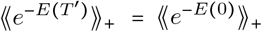, so that

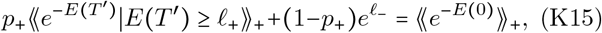

where *p*_+_ is the probability of crossing the positive threshold before crossing the negative threshold. We have used that 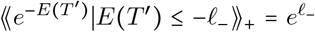 as *E* (*t*) is a continuously decreasing function. To find the survival probability, we take the limit *ℓ*_+_ → ∞ for which 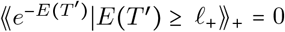 . Furthermore, setting *ℓ*_−_ = 0 and *E*(0) = *x* + Δwe find

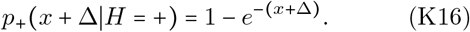

Using Eq. (K16) and Eq. (K13) in Eq. (K12), integrating over *x*, and identifying *ϵ* / *τ* =Δ/ *τ*_r_, we obtain Eq. (K3) that we were meant to derive.

**Proof for** *H* = −. We now derive Eq. (K4). The last passage time *T*_am_ for *H*= − is the largest time at which *E*(*t*) ≥ 0. Since the process *E*(*t*) decreases as a deterministic, linear function with constant slope − *ϵ* / *τ*, the last passage time takes the form

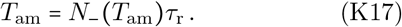

It follows that

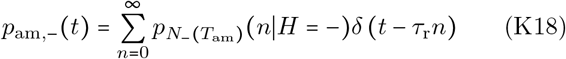

where 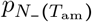 is the probability mass function of *N*_−_ *T*_am_ . Since *N*_−_ is a counting process with rate *j*_−_, the number of particles that arrives in a time interval *t = nτ*_r_ is proportional to a Poisson distribution with mean *j*_−_*τ*_r_*n*, and thus

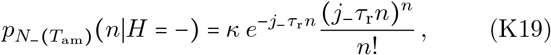

where we used the convention that 0^0^ 1, and with *κ* the proportionality constant we are left to determine. We note that the mean of the Poisson distribution depends itself on *n*. Using that 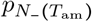 should be a normalized distribution, we find that *κ* = 1 − *j*_−_*τ*_r_ for *j*_−_*τ*_r_ < 1, which holds by the assumption *j*_+_ > *j*_−_. Substituting Eq. (K19) into Eq. (K18) yields Eq. (K4), as anticipated.

## Appendix L: Bicoid-Hunchback model

We describe the Bicoid concentration gradient as

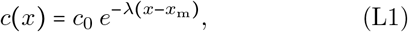

see [8, 45, 47], where *x* ∈ [0, 1] denotes the relative position along the anterior-posterior axis (*x* = 0 is the anterior pole and *x* = 1 the posterior pole, corresponding to about 500*µm*), *λ* = 5 is the inverse of the decay length of the concentration profile and *c*_0_ =4.8 ± 0.6*µm*^−3^ is the Bicoid concentration at the mid point *x*_*m*_ where the Hunchback profile switches from high to low (*x*_*m*_ ≈ 0.45 for the 10-th nuclear cycle [8, 48]). We estimate the distance between neighboring nuclei as Δ*x*= 0.02.

Due to the reproducibility of the midpoint *x*_*m*_ across embryos, and due to the sharp decline of the Hunchback expression profile at the midpoint, we assume that the two hypotheses correspond to two Bicoid concentrations close to the Hunchback boundary, separated by one nucleus diameter: *c*_+_ = *c*(*x*_m_) and *c*_−_ = *c*(*x*_m_ + Δ*x*). This choice corresponds to a precision *ϵ* = (*c*_+_ − *c*_−_)/*c*_+_ = 0.1 and a timescale *τ* = 20 s. Our model is then expressed by

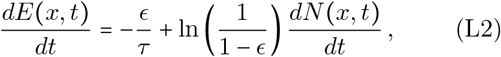

see Eq. (5), where *N* (*x, t*) is a Poisson point process in *t* of rate *j*(*x*) = 4*πrDc*(*x*), representing the rate at which Bicoid molecules hit the Hunchback promoter at position *x*. Following Ref. [47], we take *D*= 0.3(*µm*) ^2^ /*s* (but see [79]) and *r* = 3nm.

We consider the model dynamics up to a finite time *t*_f_ = 5.5 min, representing the time that a discrete nuclear envelope is observed during nuclear cycle 10, see Table 1 in Ref. [80]. The reason is that the nuclear envelope breaks down, so that the Bicoid concentration drastically decreases, at the end of each cycle [47].

**TABLE 1.**
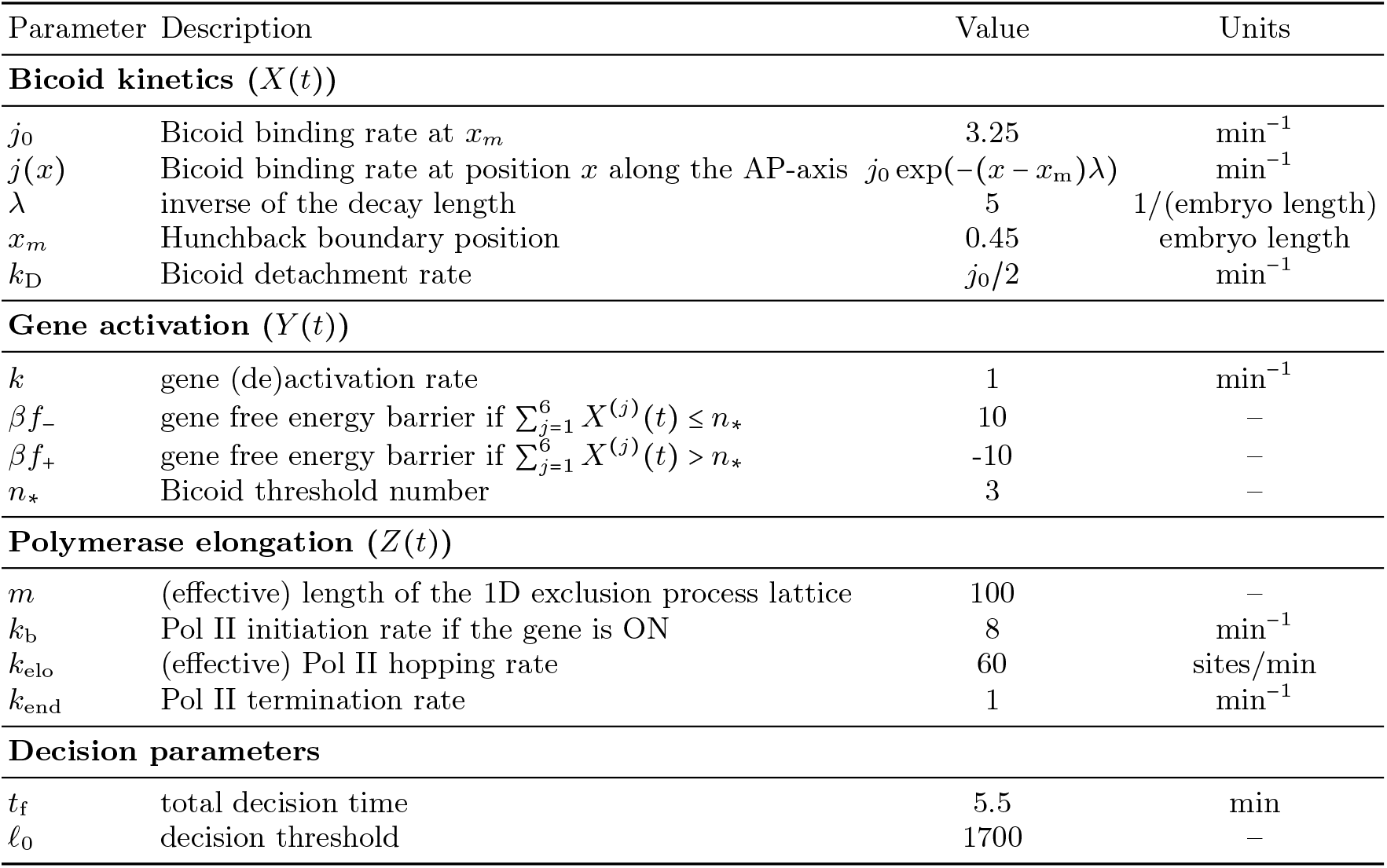
Parameters for the biophysical model for transcription (Bicoid-Hunchback system)

In the amendable model, the probability *p*_Hb_ to express Hunchback is equivalent to the probability that the evidence process is positive at time *t*_f_ . Using that *N*(*x, t*) is a Poisson random variable, we obtain

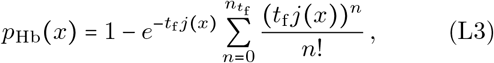

with 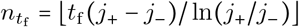 and ⌊·⌋representing the floor function.

In the irrevocable model, the final decision is taken to be + (expressing Hunchback) if the positive barrier *E*(*t*) = *ℓ*_+_ is reached by the final time, and before the *E*(*t*) reaches the negative barrier −*ℓ*_−_. The final decision is− if either the negative barrier *E* (*t*) = −*ℓ*_−_ is reached by the final time and before *E*(*t*) reaches *ℓ*_+_, or neither barrier is reached by the final time. The values of two barriers are free parameters of the decision model. We fitted these parameters to the decision profile, i.e., we minimized the mean squared distance 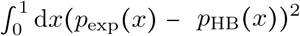 between the theoretical (*p*_HB_ *x*))^2^ and empirical(*p*_exp_ *x*)) Hunchback expression profile. This fit yields *ℓ*_+_ ≈ 0.07 and *ℓ*_−_ → ∞, see Fig. 12. This means that it is optimal to avoid committing to the negative hypothesis before the final time.

**FIG. 12.**
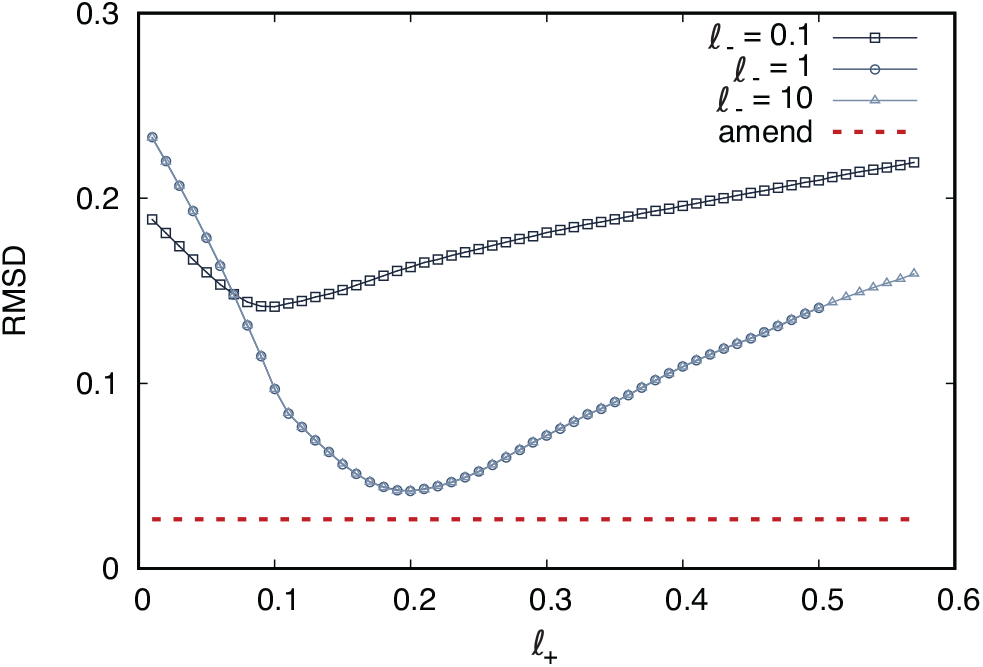
Behavior of the root mean squared deviation 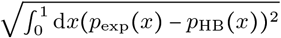 RMSD) between the predictions of the irrevocable model and the experimental Hunchback gene activation profile from Ref. [48]. The RMSD of the amendable model is shown with a red dashed line for comparison. For the irrevocable model, the behavior for *ℓ*_−_ = 1 and *ℓ*_−_ = 10 is nearly indistinguishable, signaling that the error is minimized when *ℓ*_−_ is sufficiently large. In contrast, *ℓ*_+_ needs to be tuned for minimizing the RMSD.

## Appendix M: Hunchback-Bicoid system: additional experimental comparison

We compare the theoretically predicted probability that Hunchback is expressed with experimental measurements from Ref. [48] in Fig. 8. We here provide additional information on this comparison, and also discuss different experiments.

The experiment of Ref. [48] detects Hunchback RNA transcripts that are localised to nuclei using fluorescent in situ hybridization (RNA-FISH). The authors find either no, one, or two signals per nuclei, corresponding with absence of transcription of Hunchback, active transcription in one allele, and active transcription in both alleles. Cells in the experiment are fixated, and thus this method does not provide temporal information.

In Fig. 13 we compare experimental data for the probability of Hunchback expression (purple lines of Fig. 3 in Ref. [48]) in the four nuclear cycles 10 to 13 with theoretical predictions based on the optimal amendable Bicoid-Hunchback model. The model performance remains extremely good across all four nuclear cycles. The only discrepancy is that near the anterior poles (*x* < 0.3), the Hunchback profiles in the data of Ref. [48] are substantially lower than *p*_HB_ = 1 in nuclear cycles 12 and especially 13. This effect is attributed to the activation of Torso [81], a receptor tyrosine kinase that deactivates the transcription of Hunchback near the anterior pole. The effect of this kinase is not included in our model for simplicity.

**FIG. 13.**
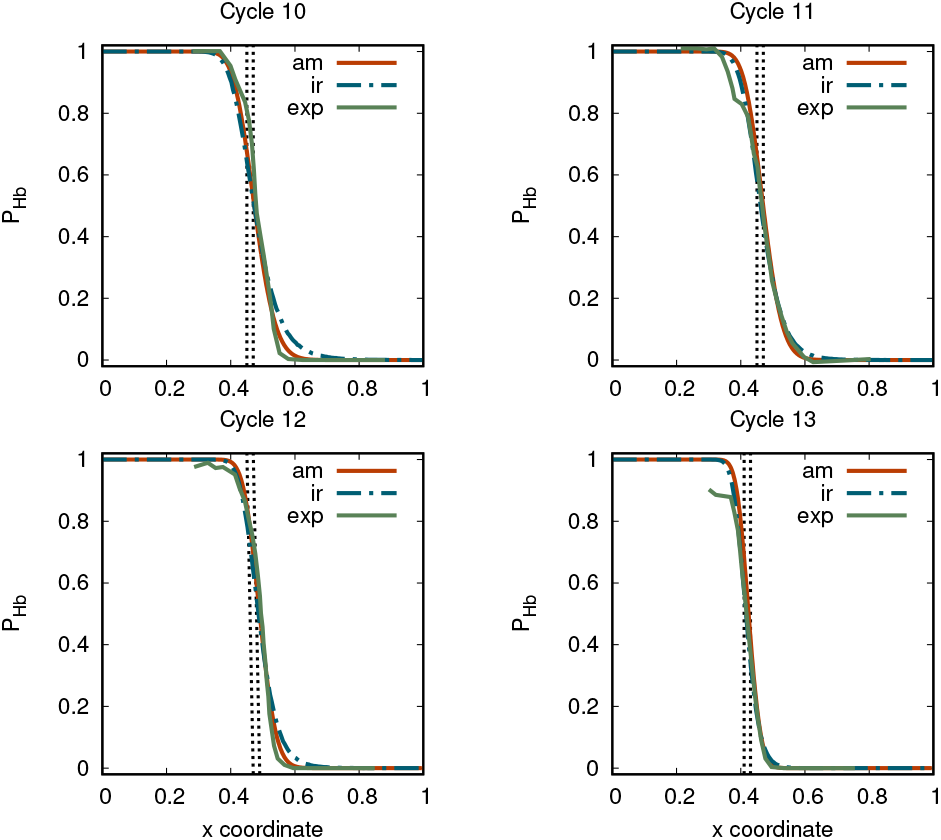
Probabilities *p*_Hb_ *x* that nuclei in the fruit fly embryo express Hunchback as a function of the relative position *x* along the anterior-posterior axis in the four nuclear cycles before cellularization. Figures compare experimental measured profiles (green solid line) with theoretical predictions based on amendable decisions (red solid line) and irrevocable decisions (blue dashed-dotted line). The curves are analogous to those in Fig. 8b, but for the nuclear cycles 10 to 13. In the nuclear cycle 14 the nuclei cellularize, at which point the expression changes significantly. In the models, the total width *t*_f_ of the time interval in which the Bicoid concentration is being sensed by the Hunchback promoter are set to 5.5 min, 6.5 min, 9.1 min, and 16 min for nuclear cycles 10, 11, 12, and 13, respectively. These estimates correspond to the time intervals for which a discrete nuclear envelope is observed, see Table 1 of Ref. [80]. All other parameters are the same for all the nuclear cycles, except the Hunchback expression mid point *xm*, which is set to 0.45, 0.45, 0.43, and 0.41 for nuclear cycles 10, 11, 12, and 13, respectively. Experimental data are the purple lines from Fig. 3 of Ref. [48].

Refs. [50–52] performed live imaging measurements of the activity of nascent Hunchback transcription in transgenic fruit fly embryos using the MS2-MCP imaging system [82]. Although Ref. [51] does not directly report the mean amendable decision times as in Fig. 8c, we shall argue that the qualitative results are in correspondence with the amendable decision model.

To perform such comparison, we first note that the mutant used in [51] expresses Hunchback in a different way than wild type embryos. Notably, in nuclear cycles 10 to 12, there is no clear anterior-posterior patterning of Hunchback (see Panels A-H in Fig. 2 of Ref. [51]). On the other hand, in nuclear cycle 13 the anterior-posterior patterning of Hunchback is present, although it presents differences from the wild type (10% of the nuclei in the mutant also express Hunchback in the posterior (right) part of the egg, in contrast with the wild type). We therefore focus on nuclear cycle 13. Figure 3C of Ref. [51] demonstrates that the time of persistent expression of nascent Hunchback transcripts in the mutant is about 8 minutes in the anterior axis (*x* < 4) and about 3 minutes in the posterior (*x >* 0.4). This supports that decisions are amendable in the posterior part of the egg. Figure 3A of [51] shows that in nuclear cycle 13 the initiation time reaches a peak around *x* ≈ 0.4, supporting our prediction of a longer decision times near the decision boundary.

## Appendix N: Amendable transcription model

The model consists of three coupled Markov jump processes: (a) an observation process *X* (*t*), representing the binding kinetics of Bicoid to the Hunchback promoter; (b) a two-state model *Y* (*t*) representing the activation-deactivation of the Hunchback allele based on the binding state of the promoter; (c) a totally asymmetric exclusion process *Z* (*t*) representing the elongation process of PolII along the allele.

We define the process *X*(*t*) closely following similar models in the literature [23, 53]. The promoter region consists of six binding sites for the transcription factor Bicoid [83–86]. Each binding site can be bound or unbound to Bicoid, so that

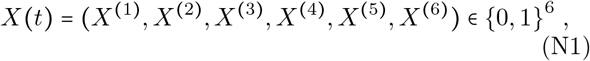

where *X*^(*i*)^ (*t*) = 1 or *X*^(*i*)^ (*t*) = 0 if the *i*-th binding site at time *t* is bound or unbound to Bicoid, respectively. If *X*^(*i*)^(*t*) = 0, then Bicoid proteins can bind to the *i*-th binding site at a rate *j*(*x*) = 4*πrDc*(*x*), see Appendix L. If *X*^(*i*)^(*t*) = 1, then the Bicoid bound at the *i*-th site can detach at a rate *k*_D_.

We represent the gene state by a single variable *Y* (*t*) *∈* {0, 1} with *Y* (*t*) = 0 if the gene is in the OFF state and *Y* (*t*) = 1 if the gene is in the ON state. The gene switches between the two states at rates

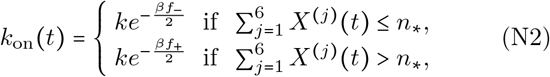

and

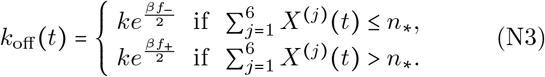

Here *n*_*\**_, *k, βf*_−_, and *βf*_+_ are model parameters. We choose *f*_−_ > 0 and *f*_+_ < 0 so that the gene is more likely to switch ON if the number of bound Bicoid transcription factors exceeds *n*^*\**^, while the gene is more likely to switch OFF if the number of bound Bicoid transcription factors is smaller or equal than *n*_*\**_ . The parameters *f*_−_ and *f*_+_ can be interpreted as free energy differences between the ON and the OFF state and *β* as the inverse temperature.

The last component of the model is the actual transcription process, which we model as a variant of an exclusion process [54]. The transcribed region of the DNA is modeled as a discrete lattice with *m* sites. Each site can be occupied by at most one polymerase. Therefore, the state of the lattice at time *t* is described by

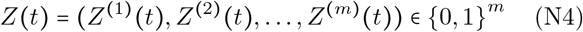

with *Z*^(*i*)^(*t*) = 1 if the *i*-th site is occupied by a polymerase, and *Z*^(*i*)^(*t*) = 0 if the *i*-th site is unoccupied. Changes in *Z*(*t*) are determined by the kinetics of the polymerases, which depend on *Y* (*t*). Polymerases hop forward at a rate *k*_elo_ regardless of the value of *Y* (*t*), and they detach from the DNA at a rate *k*_end_ when they have reached the last, *m*-th site. If the gene is ON, then a polymerase can bind to the first site of the lattice at a rate *k*_b_, thereby initiating transcription.

Model parameters are summarized in Table 1. The rate *j*(*x*) is the same as in Section VII, see also Appendix L. The parameter *k*_D_ is experimentally unknown. However, as the decision profile changes around the midpoint, we require that the number of bound transcription factors approximately equals *n*_*\**_ near the midpoint, and therefore we set 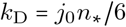. We set the parameter *k* = 1/min, following Ref. [52] that infers the timescales between gene switching events from experimental data. The parameters *n*_*\**_, *βf*_−_, and *βf*_+_ are not known, to the best of our knowledge. We set *n*_*\**_ = 3, *βf*_*_*_ − /2 = 5 and *βf*_*+*_ /2 = −5. The rate of initiation of polymerases is about *k*_b_ = 8/ min [52]. The elongation rate is 1.8kb/min [52] and the length of the transcript is 3kb [87]. However, as polymerases occupy about 30 nucleotides, we set *m* 100 and the elongation rate is thus *k* = 60 sites/min. We set the termination rate equal to *k*_end_ = 1 /min [52], allowing some time for the transcript to maturate before detaching. The total run time is taken to be *t*_f_ = 5.5min, corresponding with the time allowed for transcription in the 10th nuclear cycle [80].

### 1. Biophysical decision model

We now define the evidence and the decision processes for the amendable transcription model. We take as the evidence process the sum of the lengths of the RNA transcripts bound to PolII minus a constant threshold *ℓ*_0_,

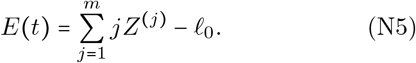

The decision variable *D* = sign(*E*(*t*_f_)) denotes whether Hunchback is transcribed (*D* = +) or not (*D* = −).

The choices of the evidence process and the decision variable are motivated by the experiment of Ref. [48], that measured the presence of nascent RNA transcript bound to the DNA. Hence *D* = + corresponds with the measured presence of RNA transcript localised to the DNA, and thus presumably bound to PolII. We subtract a constant *ℓ*_*0*_ from the total length 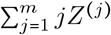 to account for the fact that the RNA transcript must be large enough to be experimentally detectable. We have set *ℓ*_0_ = 17*m* as this yields the sharpest decision profile in our model.

For the optimal decision model, we consider the evidence process [23, 28]

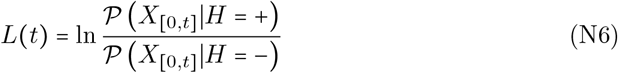

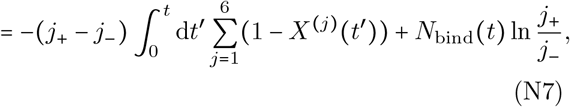

where *N*_bind_(*t*) is the number of Bicoid binding events in the time interval [0, *t*]. Considering that nuclei are separated by a distance Δ*x* = 0.02, we have set the rates for the two hypotheses equal to *j*_+_ = *j* (0.46) and *j*_−_ = *j* (0.48) . We remark that this optimal decision model is more detailed than the one presented in Section VII, in that it assumes a finite unbinding rate and explicitly considers the presence of six binding sites for Bicoid. However, the two optimal models yield very similar predictions, see Fig. 9b.

Trajectories of the evidence process associated with the amendable transcription model illustrate how the system continuously updates the evidence based on the amount of bound Bicoid, see Fig. 14. In the figure we also show the corresponding optimal evidence processes *L*. We note that *E* is a suboptimal evidence process and is thus not trivially linked with *L*. However, when *E* is large, then also *L* tends to be large, see for example the purple curves for *x* = 0.5.

**FIG. 14.**
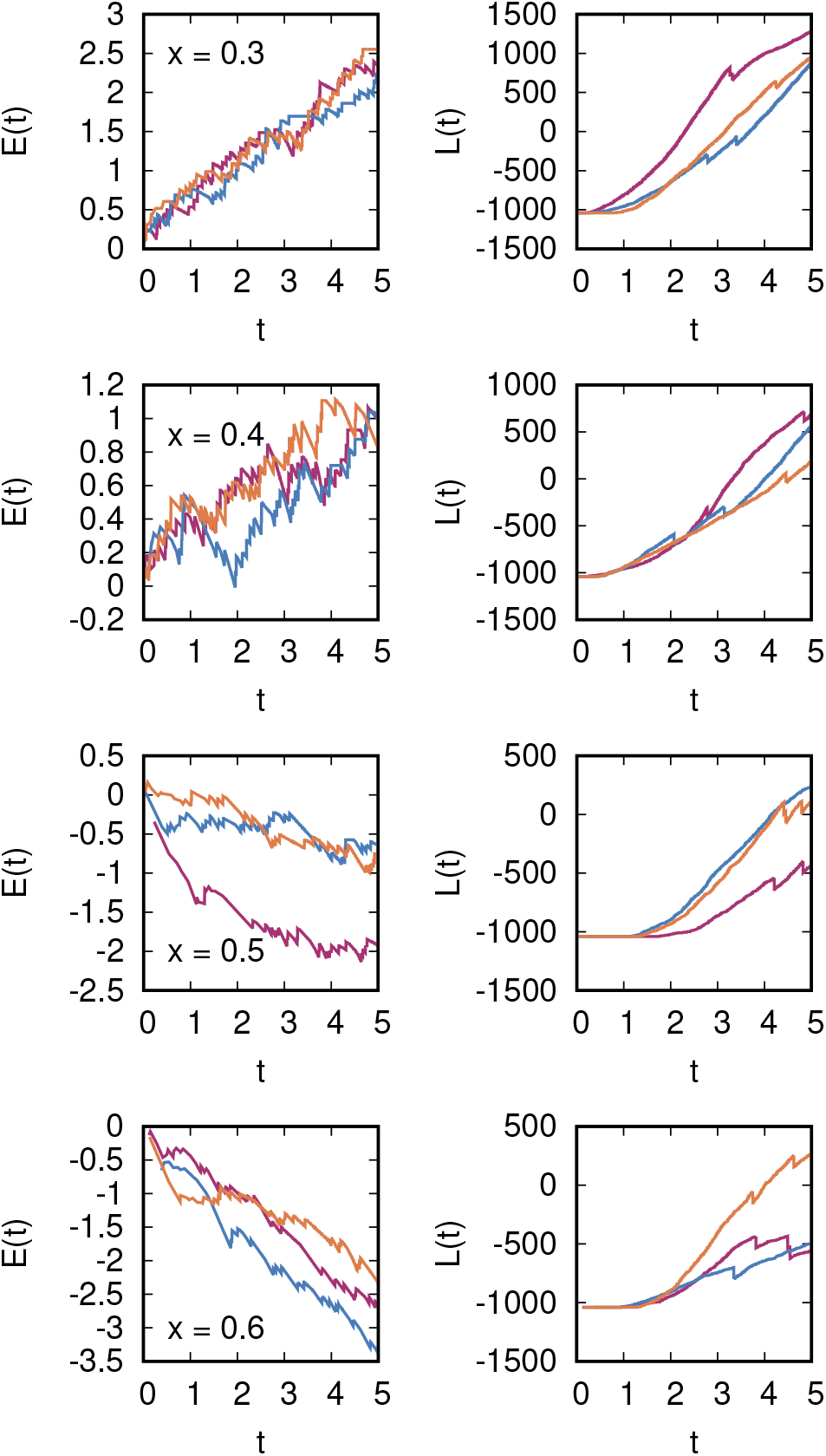
(left) Sample trajectories of the evidence process *E* (*t*) for the biophysical transcription model, see Eq. (N5), for relative coordinate values *x* = 0.3, *x* = 0.4, and *x* = 0.6 along the anterior-posterior axis. For other model parameters, see Appendix N. The three colors represent three different independent realizations of the process for each value of *x*. (right) Corresponding optimal evidence processes *L*(*t*), as given by Eq. (N7), evaluated for the same realizations of *X*(*t*).

